# Transient telomere uncapping triggers telomeric and subtelomeric rearrangements

**DOI:** 10.1101/2025.07.18.665472

**Authors:** Liébaut Dudragne, Clotilde Garrido, Oana Ilioaia, Juliana Silva Bernardes, Zhou Xu

## Abstract

Telomeres are nucleoprotein structures that cap the extremities of eukaryotic linear chromosomes, thus preventing them from being detected as DNA damage. Telomere uncapping poses a profound threat to genome integrity, yet the immediate consequences of transient uncapping remain unclear. In *Saccharomyces cerevisiae*, the Cdc13-Stn1-Ten1 complex caps telomeres and limits resection, which would otherwise lead to DNA damage checkpoint activation. Here, using the temperature-sensitive *cdc13-1* allele, we demonstrate that even transient telomere uncapping induces extensive genomic rearrangements within a few cell cycles, despite a functional DNA damage checkpoint. Two distinct rearrangement signatures were observed in cells surviving transient uncapping: one characterized by the reorganization and recombination of the subtelomeric region, mostly involving the Y’ elements, and the other exhibiting massively elongated telomeres up to 10 kb, corresponding to a ∼30-fold increase. Long-read sequencing revealed that the genomic instability was confined to the subtelomere and telomere regions, and evidenced Yʹ element loss/amplification, terminal duplication of chromosome ends, and telomeric-circle-driven amplification of telomere repeats. Rearrangements unfold over multiple generations and require the homologous recombination factor Rad52 and the Polδ subunit Pol32, which is essential for break-induced replication. The recombination proteins Rad51 and Rad59 also contribute to the rearrangements in partially independent pathways. Remarkably, survivors with elongated telomeres demonstrate robust resistance to subsequent telomere uncapping, in a Rad52-dependent manner. Our findings provide novel insights into the consequences of transient telomere uncapping for genome stability, a process that might contribute to subtelomere and telomere dynamics and evolution.

## Introduction

Protection of chromosome ends is critical for genome integrity. In most Eukaryotes, it is ensured by specialized repetitive DNA sequences, known as telomeres, that prevent chromosome ends from being recognized and processed as DNA damage (de Lange, 2018; Jain & Cooper, 2010). However, telomeres shorten at each cell division due to the end-replication problem. This process is compensated for by the recruitment of telomerase, an enzyme capable of elongating telomeres and maintaining them at a steady-state length distribution (Wellinger & Zakian, 2012). In *Saccharomyces cerevisiae*, telomeric sequences are composed of ∼350 bp of degenerated TG_1-3_ repeats with a short (5-15 nt) 3’ single-stranded overhang (Larrivée et al., 2004; Soudet et al., 2014).

In yeast cells lacking telomerase, telomeres continue shorten to eventually reach a critical length at which point cells activate the DNA damage checkpoint and stop dividing, in a process called replicative senescence (Enomoto et al., 2002; IJpma & Greider, 2003; Lundblad & Szostak, 1989). Genome instability, especially near chromosome ends, has been shown to increase during this phase (Coutelier et al., 2018; Hackett et al., 2001; Hackett & Greider, 2003). In prolonged telomerase-negative cultures, rare cells can escape senescence at an estimated frequency of ∼10^-5^ per generation by engaging in alternative, recombination-based pathways to maintain their chromosome ends (Chen et al., 2001; Kockler et al., 2021; Le et al., 1999; Lundblad & Blackburn, 1993; Teng et al., 2000; Teng & Zakian, 1999). These post-senescence survivors emerge through two distinct homology-dependent mechanisms: type I survivors amplify the subtelomeric Y’ elements, whereas type II survivors elongate telomeric repeats, a mechanism also found in ALT cancer cells (Cesare & Reddel, 2010). Both types of survivors depend on Rad52, which is required for all types of recombination in yeast, and Pol32, a subunit of polymerase δ involved in break-induced replication (BIR) (Lundblad & Blackburn, 1993; Lydeard et al., 2007). Other genetic requirements for both types have been investigated in depth. For example, type I and type II survivors depend on the recombination factors Rad51 and Rad59, respectively (Chen et al., 2001; Teng et al., 2000). Although the type I and type II pathways have long been thought to be independent, a recent work proposed a unified pathway in which Rad51 is required for the formation of precursors, followed by Rad51-independent but Rad59-dependent maturation into stable survivors (Kockler et al., 2021). Thus, telomere shortening leads to genome instability and rearrangements in telomerase-independent survivors, which in turn bypass the need for telomerase to maintain telomeres.

Telomere instability can also be induced when telomere protection fails. Telomeres are normally capped by the binding of specific proteins. Defects in these telomere-capping proteins or their binding have been shown to cause inappropriate repair events, such as telomere fusions or recombination involving telomeric or subtelomeric sequences (Fellerhoff et al., 2000; Garvik et al., 1995; Grandin, 2001; Grandin & Charbonneau, 2003; Marcand et al., 2008; Mieczkowski et al., 2003; Zubko & Lydall, 2006). These proteins include Rap1 and its cofactors Rif1, Rif2 and the Sir complex, the Ku complex, as well as the CST (Cdc13-Stn1-Ten1) complex (Wellinger & Zakian, 2012). Cdc13 is a key telomeric factor given its triple role in telomere maintenance: it assists telomere replication together with Stn1 and Ten1, promotes telomere elongation by recruiting telomerase and prevents excessive degradation of the 5’ end by nucleases (Churikov et al., 2013). The *cdc13-1* allele is a temperature-sensitive mutant widely used to study telomere capping (Garvik et al., 1995; Hartwell & Smith, 1985; Mersaoui & Wellinger, 2019). The incubation of the *cdc13-1* mutant at restrictive temperatures (≥ 32°C) leads to resection of the 5’ end and accumulation of single-stranded DNA (ssDNA) at telomeres, which activates the DNA damage checkpoint (Garvik et al., 1995; Lydall & Weinert, 1995). Permanent incubation of *cdc13-1* cells at restrictive temperature leads to cell death, despite a limited number of cell divisions allowed by adapting to the checkpoint in the first 24 hours (Lee et al., 1998; Sandell & Zakian, 1993; Toczyski et al., 1997). Cells resistant to permanent telomere uncapping can only be selected if the resection or checkpoint pathways are also altered (Grandin, 2001; Grandin & Charbonneau, 2003, 2013; Zubko & Lydall, 2006), suggesting that cells with deprotected telomeres are eliminated in conditions with functional DNA processing and checkpoint, thus safeguarding genome integrity. For example, checkpoint-deficient *mec3Δ cdc13-1* cells incubated at an intermediate temperature of 29°C tolerated telomere deprotection and, after several passages, generated Cdc13-independent survivors with very long telomeres akin to those of telomerase-negative type II survivors (Grandin, 2001). Similar survivors were also generated in *cdc13-1 exo1Δ* mutants grown at 36°C, and at a higher frequency in the checkpoint-deficient *cdc13-1 exo1Δ rad9Δ* mutant (Zubko & Lydall, 2006).

Overall, Cdc13 dysfunction alone, without additional mutations in checkpoint- or resection-related genes, has not been shown to lead to the emergence of cells with rearranged telomere structures. In addition, previous studies of the long-term consequences of permanent telomere deprotection might have missed the direct molecular response to telomere deprotection. Here, instead, we investigated the early effects of telomere deprotection on genome stability in the presence of a functional checkpoint. To do so, we use an experimental protocol where telomeres are transiently deprotected and cells are allowed to recover, thus enabling us to characterize the full spectrum of molecular consequences, including events that would have been counterselected if telomeres were kept deprotected permanently. We find that genomic instability arises earlier than anticipated, even in the presence of a functional checkpoint, and is limited to the telomere and subtelomere regions. Using long-read sequencing and telomere-to-telomere genome assembly, we characterized Rad52- and Pol32-dependent rearrangements of the subtelomeric Y’ elements and telomere elongation up to 8-10 kb. Strikingly, we discovered that very long telomeres together with Rad52 protect cells from a second telomere uncapping event, suggesting that this telomeric nucleoproteic structure acts as a Cdc13-independent protective cap. Our results provide new insights into telomere-driven genomic instability and underscore the impact of transient telomere dysfunction on chromosome integrity.

## Results

### Transient telomere uncapping impairs cell survival

To evaluate the direct molecular and cellular response to telomeric uncapping, we employed the temperature-sensitive *cdc13-1* allele to induce a transient uncapping. We exposed the *cdc13-1* strain to the restrictive temperature (32°C) for various durations. The cells were first grown overnight in rich liquid medium at the permissive temperature (23°C), then plated on solid YPD media and incubated at 32°C for a defined time *“t”* (0–72 h) to induce telomere uncapping (Fig. 1A). The cells were then returned to the permissive temperature to assess recovery and survival. The cells tolerated uncapping for up to 6 h without a noticeable decrease in viability, but the survival frequency decreased significantly for *t* ≥ 12 h (Fig. 1B and C). Some rare colonies formed even after uninterrupted incubation at 32°C for 72 h, establishing a lower limit for cell survival to *cdc13-1*-induced telomeric uncapping at 32°C. The survival frequency for *t* = 24 h was 5.5-fold greater than for *t* = 72 h (Fig. 1C).

**Figure 1.**
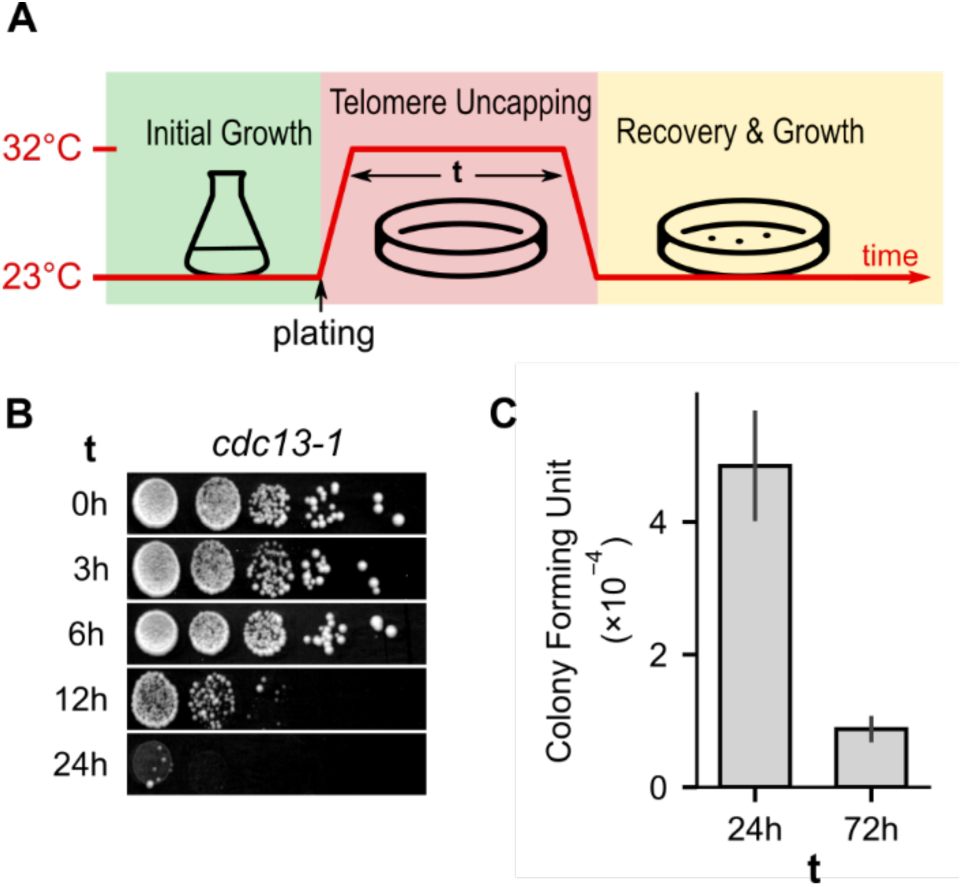
Effect of transient telomere uncapping on survival. (A) Scheme of the experimental protocol: *cdc13-1* cells were first grown at permissive temperature (23°C) before plating and incubation at restrictive temperature (32°C) for a duration *t*. The plate was then returned to 23°C for recovery and growth. Individual colonies were picked for further analysis. (B) Spot assay showing the growth of *cdc13-1* cells on YPD rich media after *t* hours of telomere uncapping. (C) Colony formation frequency measured using the protocol shown in (A). A control plate was maintained at 23°C for normalization. The error bars correspond to the standard error of the mean; n = 7 (24 h) and n = 4 (72 h) independent experiments.

We conclude that transient Cdc13 dysfunction at telomeres impairs cell survival in a time-dependent manner. To investigate the molecular changes induced by transient uncapping, we selected the 24-h-long uncapping condition for further analysis.

### Cells exposed to telomere uncapping for 24 h show extensive genomic rearrangements

To study the impact of transient *cdc13-1*-mediated uncapping on genomic stability, we selected and grew colonies formed from cells that survived 24 h at 32°C. We investigated large-scale genomic rearrangements in these survivors using Pulsed Field Gel Electrophoresis (PFGE), which enables the resolution of whole chromosomes on agarose gels. Among the 7 analyzed colonies, 6 presented significant shifts in chromosome length, corresponding to changes of at least tens of thousands of base pairs (Fig. 2A).

**Figure 2.**
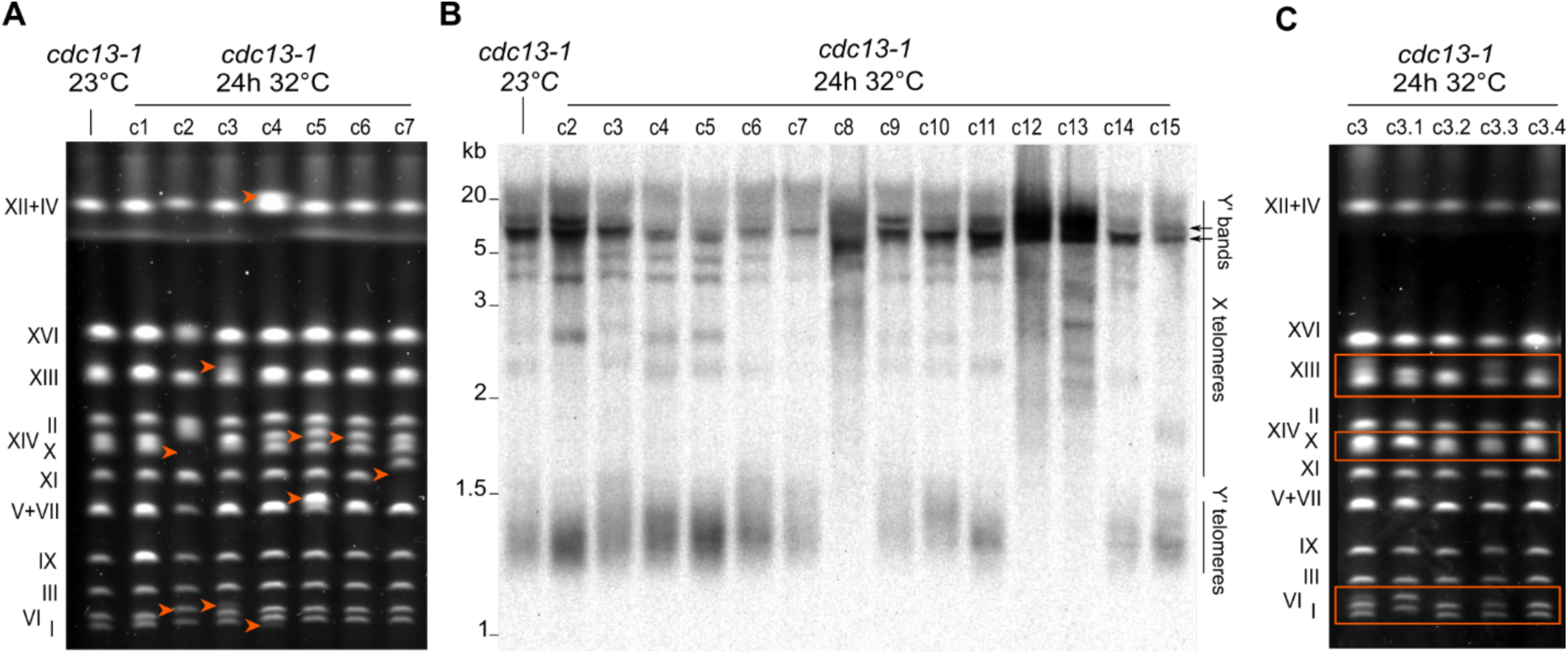
Transient telomere uncapping leads to chromosomal instability and changes in telomere and subtelomere structure. (A) PFGE of 7 *cdc13-1* survivor clones as well as a *cdc13-1* control strain grown constantly at 23°C. Compared to the control strain, 6 out of the 7 survivor clones exhibited apparent chromosome size shifts, marked by orange arrows. (B) TRF Southern blot analysis of 14 *cdc13-1* survivor clones and a control strain. The genomic DNAs were cut with the *XhoI* enzyme and a telomeric probe was used. (C) PFGE of 4 subclones of clone 3 from (A) and (B), derived from single cells grown at 23°C. The orange boxes indicate bands with distinct migration patterns in the 4 subclones.

Regions adjacent to the telomeric repeats typically contain Telomere-Associated Sequences (TASs), namely X and Y’ elements. Y’ elements, present in zero, one, or multiple tandem copies separated by short interstitial telomeric sequences, exist in two main size classes: long (6.7 kb) and short (5.2 kb). The X element, which is found at all extremities in laboratory strains, varies in sequence and size. To test whether telomeres or TASs are involved in the observed instability, we employed Terminal Restriction Fragment (TRF) Southern blotting, which probes telomere sequences in *XhoI*-restricted fragments to inform on characteristic features of the telomere structure. In the control *cdc13-1* strain grown at 23°C, telomeres associated with Y’-containing subtelomeres (“Y’ telomeres” in Fig. 2B) migrated to form a smear near 1.2 kb, corresponding to ∼300 bp of telomeric repeats and a ∼900 bp segment defined by the *XhoI* site in the Y’ element. In contrast, telomeres at X-only subtelomeres (’X telomeres’ in Fig. 2B) migrated as longer fragments owing to more distal *XhoI* restriction sites. Additionally, interstitial telomere sequences between Y’ elements allowed probing of tandem Y’ repeats (“Y’ bands” in Fig. 2B). Out of 31 clones that survived transient telomere uncapping, 6 exhibited very long and heterogeneous telomeres (c8, c12, c13, c20, c21, and c27, in Fig. 2B and Supp. Fig. S1A), a pattern that resembles that observed in telomerase-independent, recombination-dependent, type II survivors (Lundblad & Blackburn, 1993). Almost all other survivors showed visible alterations in Yʹ-specific-band intensity and/or loss of X-associated telomeric bands, indicative of putative Y’ amplification or acquisition at X-only subtelomeres, respectively. Y’ amplification and acquisition are also observed in type I post-senescent survivors, although to a much greater extent where no X band is retained and the Y’ signal is strongly amplified. Notably, we observed no telomere shortening in any clone, which is consistent with telomerase being active (Fig. 2B). We referred to these uncapping-induced rearrangement patterns as type-II-like (T-II-L) survivors for the survivors with lengthened telomeres and Y’-associated survivors (YAS) for those only exhibiting altered Y’ organization.

### Genomic instability unravels over multiple cell divisions independently of checkpoint adaptation

The PFGE profiles of several clones presented blurred bands or bands of different intensities, suggesting intraclonal heterogeneity (Fig. 2A). To test this possibility, 3 primary clones were subcloned, and 4 subclones of each were reanalyzed by PFGE. All 3 primary clones exhibited subclonal heterogeneity in their PFGE patterns, revealing that genomic instability persisted during several cell divisions (Fig. 2C and Supp. Fig. S1B). This was particularly obvious in clone c3, where no 2 subclones shared the same pattern, indicative of at least 2 rounds of divisions with ongoing rearrangements. Whether these cell divisions occurred while telomeres were still uncapped or after returning to 23°C is unclear.

Indeed, after prolonged activation (∼8 h) of the DNA damage checkpoint, cells can undergo checkpoint adaptation (Lee et al., 1998; Sandell & Zakian, 1993; Toczyski et al., 1997), resuming cell cycle despite unrepaired DNA damage and allowing for a few cell divisions while telomeres are uncapped. This process is associated with increased genomic instability (Coutelier et al., 2018; Galgoczy & Toczyski, 2001). To assess the role of checkpoint adaptation both in genomic instability and in intraclonal heterogeneity, we introduced the adaptation-deficient *cdc5-ad* allele of *CDC5*, encoding the Polo kinase essential for late cell cycle events and for adaptation (Toczyski et al., 1997). *cdc5-ad cdc13-1* cells that survived 24 h of telomere uncapping exhibited levels of genome rearrangements and intraclonal heterogeneity comparable to the *CDC5 cdc13-1* strain (Supp. Fig. S1C and D). Since *cdc5-ad cdc13-1* cells do not divide while at 32°C, this finding indicates that genomic instability could persist over several divisions after telomere protection is re-established. To circumvent this intraclonal heterogeneity, all subsequent analyses were conducted using subclones of survivor colonies.

### Genomic rearrangements are confined to telomeric regions

Given the important size shifts observed on the PFGE gels, we wondered whether the observed genomic instability was confined to the telomeric regions. To comprehensively assess the localization of the rearrangements, we performed Oxford Nanopore long-read sequencing on the genomic DNA of 9 survivors as well as the *cdc13-1* control strain, which did not experience telomere uncapping. The longest and highest quality Nanopore reads were assembled, and the assemblies were manually inspected to correct any assembly mistake in telomeric regions. This procedure yielded highly contiguous, high quality telomere-to-telomere genome assemblies (Supp. Table 2).

To investigate the rearrangements in telomeric regions, we first focused on the dynamics of Y’ elements. To track them accurately, Y’ elements identified in the control strain genome assembly were labeled and clustered based on sequence similarity (Fig. 3A). We identified 34 Yʹ elements found at 20 out of 32 extremities, with 0 to 6 tandem Y’ elements per extremity (Supp. Data 1). Among the 34 Y’ elements, we detected 20 unique sequence variants that could be grouped into 11 clusters. Two elements were classified as the same variant if their sequences were identical within the expected margin of assembly errors. Clustering analysis recovered the two main size families of Yʹ elements—short and long—along with finer sub-clusters, revealing the presence of outliers alongside large groups (Fig. 3A). Interestingly, although Y’ elements are typically classified as either 5.2 kb or 6.7 kb in size, we observed that Y’ short elements ranged from 5,062 bp to 5,592 bp (mean: 5,470 bp), Y’ long elements ranged from 6,484 bp to 6,933 bp (mean: 6,670 bp), and one element exhibited an intermediate size of 5,981 bp.

**Figure 3.**
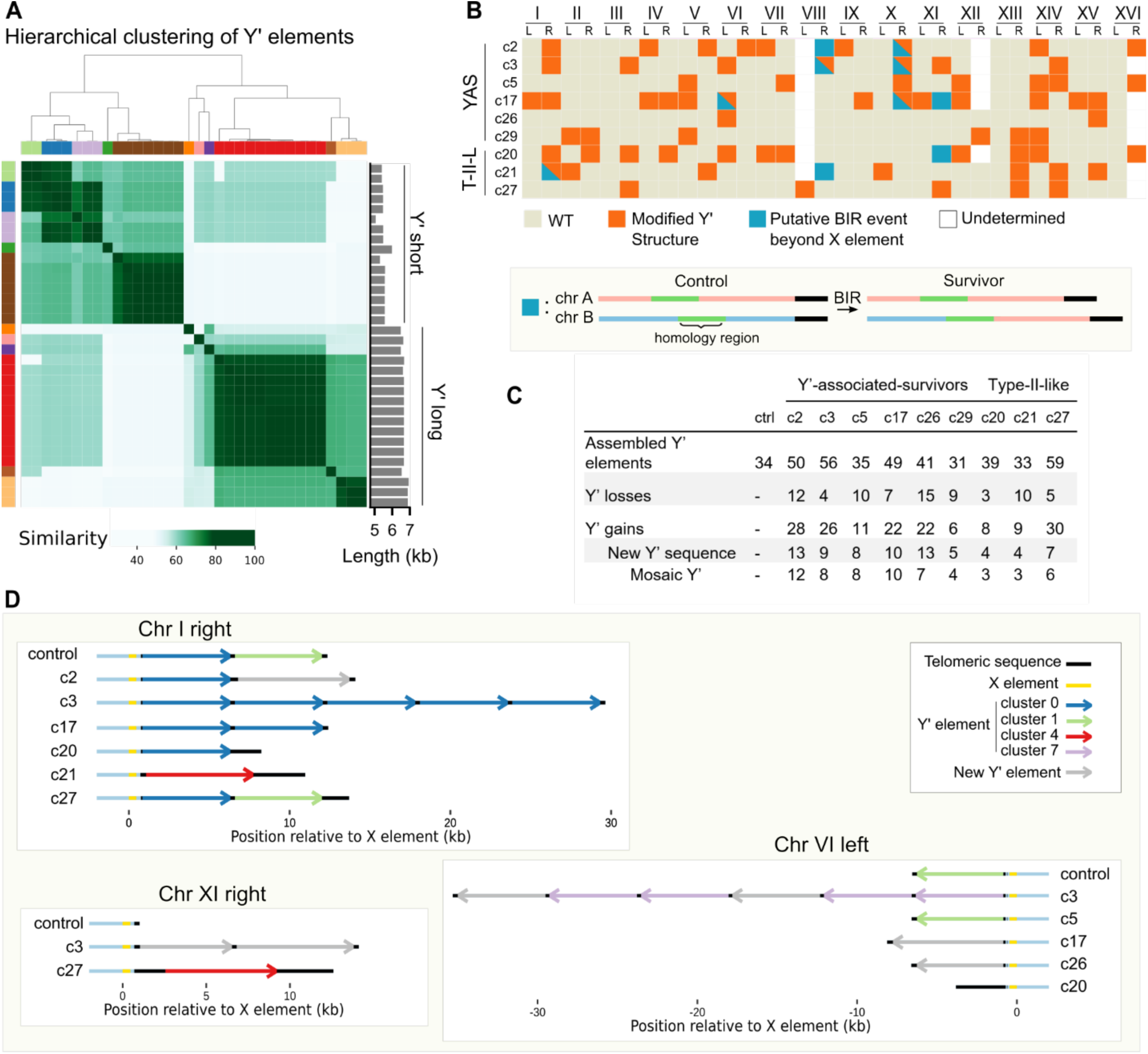
Subtelomeric Y’ elements are rearranged following telomere uncapping. (A) Similarity matrix and clustering of Y’ elements of the control strain. A similarity measure was computed for each pair of Y’ sequences and used for subsequent hierarchical clustering (dendrogram shown above the matrix). The colors correspond to the different clusters (n = 11 clusters). The bar plot on the right side indicates the corresponding lengths of the Y’ sequences. See Supp. Data 1 for the sequences of all 34 Y’ elements. (B) Map of alterations at all chromosome extremities. The labelling of Y’ elements was used to detect changes of Y’ structure at a given telomere (orange square or triangle), and duplications in core genome sequences indicate probable BIR events (blue square or triangle). YAS = Y’-associated survivors, T-II-L: Type-II-like survivors. Extremities with more than 5 Y’ elements in the reference could not always be resolved with high confidence and are thus marked as “undetermined”. (C) Summary table of the total number of indicated Y’ changes in a survivor clone. (D) Visual representation of an illustrative subset of modified Y’ structures. Three extremities are displayed as examples of typical modifications of subtelomere structure found in the indicated clones, as compared with the control: Y’ losses, gains, tandem duplications, and new Y’ sequence emergence. The long telomere sequences of type-II-like survivors (c20, c21 and c27) are apparent. See Supp. Data 2 for the representation of all extremities across all sequenced clones.

Using this labeling and clustering approach, we quantified the number of Yʹ elements associated with rearrangement events in the clones surviving transient uncapping (Fig. 3B and C). On average, 8.3 ± 3.7 (mean ± SD) Y’ elements were lost and 18 ± 8.9 were gained in each survivor clone, affecting 7.4 ± 3.1 extremities, and far exceeding what could be confidently detected by TRF Southern blot analysis. These events included loss or gain of a known Y’ element, tandem amplification of existing elements, and most interestingly, emergence of new Y’ elements whose sequences did not correspond to any element present in the control (Fig. 3C and D, Supp. Data 2). The frequency of these Y’ structure alterations was significantly higher at Y’ telomeres than at X telomeres (28% vs. 15%; p-value = 0.007, Mann-Whitney test). We found that at least 84% of the new Y’ elements could be explained as mosaics of 2 or more other Yʹ elements, suggesting that most of them originated from recombination events between existing Y’ elements rather than mutation accumulation. Interestingly, type-II-like survivors exhibited levels of Y’ rearrangements comparable to Y’-associated survivors (YAS: 7.3 ± 3.3 extremities affected, T-II-L: 7.7 ± 2.5 extremities affected; mean ± SD).

We also examined whether genome rearrangements extended beyond the X and Y’ elements into more centromere-proximal regions. We found 10 cases of terminal duplications from one chromosome end to another, spanning between 850 bp and 89 kb of sequences centromere-proximal to the X element across 5 clones (Fig. 3B). In 2 cases, 2 events were found at the same extremity, with one duplication contained within the other. By mapping these regions against the control genome, we found that these events consistently involved junctions in regions of homology between the donor and recipient chromosome arms, the size of which ranged from several hundreds to several thousands of base pairs. Based on the presence of homology and on the pattern of terminal duplication, these events likely resulted from break-induced replication (BIR).

Finally, we used the variant callers Sniffles2 (Smolka et al., 2024) and PAV (Ebert et al., 2021) to identify other structural rearrangements (>50 bp) that might have occurred in the rest of the genome. No event was detected aside from those described above, indicating that genome rearrangements were confined to telomere-proximal regions.

### Type II-like survivors exhibit tandem repeat amplification of telomeres

To study the impact of uncapping on telomere sequences, telomere length distributions were measured from the sequencing reads of all sequenced survivors (Garrido et al., 2025). In Y’-associated survivors, no significant change in telomere length distribution was observed in any clone (Fig. 4A), which is consistent with the results of TRF Southern blot analysis (Fig. 2B and Supp. Fig. S1A). In contrast, we confirmed that type-II-like strains presented very heterogeneous and long telomeres of up to 10 kb (Fig. 4B and C, and Supp. Fig. S2A-C). Telomere length distributions in type-II-like survivors presented multiple peaks, explained by the heterogeneity in chromosome extremity-specific average lengths, including a major peak aggregating 4-7 extremities with an average telomere length < 500 bp (Fig. 4C and Supp. Fig. S2B-C), indicating that not all telomeres were elongated equally.

**Figure 4.**
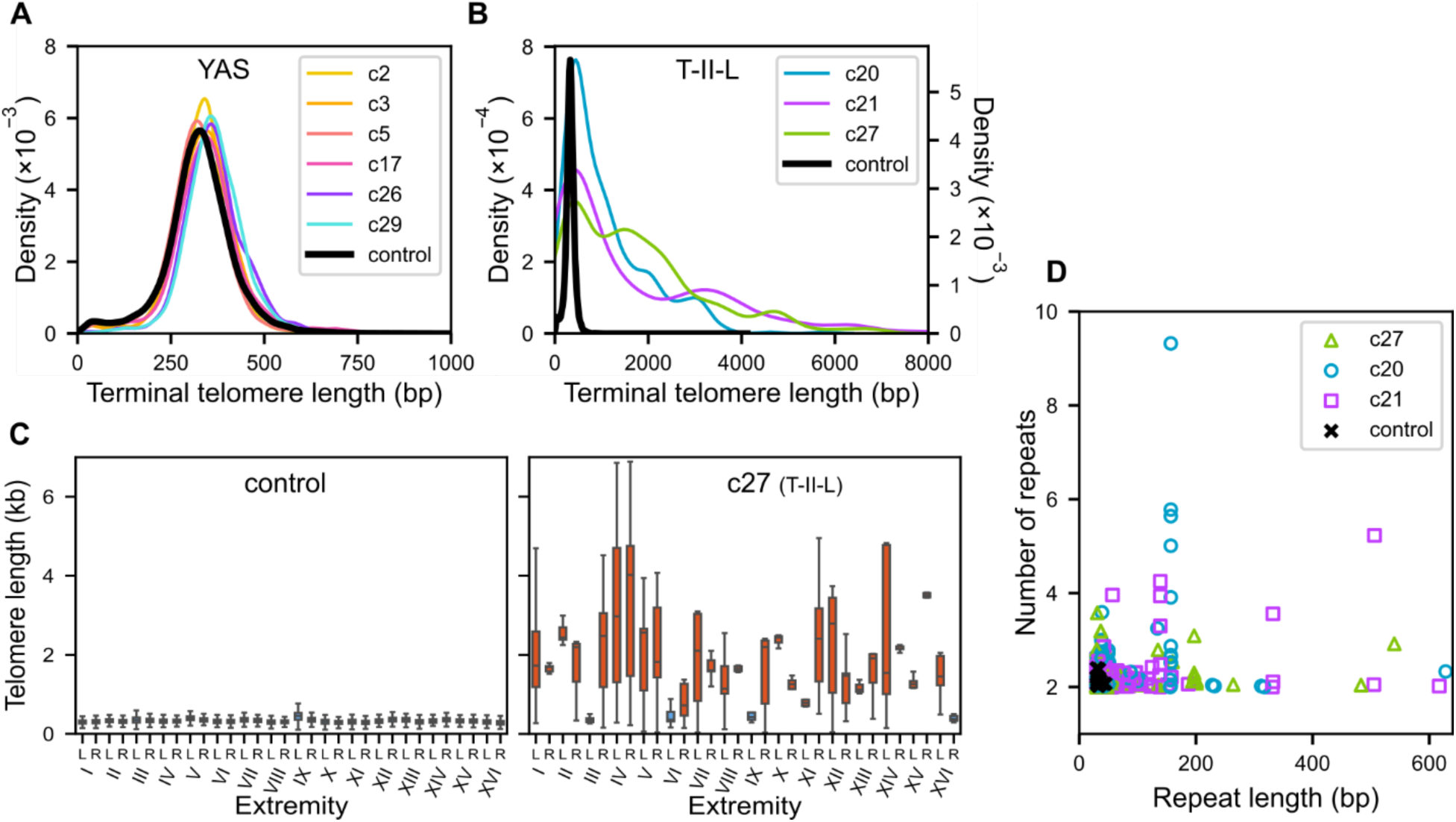
Telomeres are elongated heterogeneously in T-II-L survivors and contain large perfect tandem repeats. (A) Telomere length distributions of YAS strains compared to the control. (B) Telomere length distribution of T-II-L survivors. The left y-axis is used for survivors, whereas the right y-axis is used for the control strain. The x-axis is truncated at 8000 bp. (C) Boxplots representing the telomere length distribution at each chromosome extremity. The control strain (left) displays extremity-specific mean telomere lengths of approximately 300-400 bp with limited variations between extremities (see also Supp. Fig. S2A), whereas survivor c27 shows heterogeneously elongated telomeres with great variations between extremities (see Supp. Fig. S2B-C for other T-II-L survivors). The blue boxplots correspond to distributions with a mean telomere length < 500 bp. (D) Arrays of perfect tandem telomere repeats detected in the genome assemblies of T-II-L survivors and the control strain are represented in a 2-D plot showing the number of repeats and the length of each repeat.

Telomere lengthening in telomerase-negative post-senescence survivors is attributed to recombination events and rolling circle amplification of telomeric circles (t-circles) (Aguilera et al., 2022; Larrivée & Wellinger, 2006; Lin et al., 2005; Natarajan & McEachern, 2002). Telomerase in budding yeast adds imperfect repeats and creates a degenerated telomere motif (Forstemann, 2000). As a consequence, long enough telomere sequences generated by telomerase are unique and can be identified unambiguously. In contrast, recombination between telomeres or rolling circle amplification is expected to yield 2 or more copies of identical sequences. We thus looked for tandem amplification of telomeric sequences. In the control strain, we found a few tandem sequences of 2 copies for repeats of < 45 bp. These tandem repeats could either be signatures of previous events of recombination at telomeres or instances in which telomerase stochastically added 2 identical sequences in tandem. To specifically capture signatures of telomere uncapping, we excluded all repeats shorter than 50 bp from our analysis of type-II-like survivors. We found that the 3 sequenced type-II-like strains contained many perfect tandem repeats at telomeres. While we detected relatively short sequences (< 100 bp) that were repeated 3 times or fewer in tandem, some repeats were longer than 600 bp and others were found in arrays of up to 9 copies (Fig. 4D and Suppl. Fig. S2D). We found multiple raw Nanopore reads supporting these tandem repeats, thus excluding assembly errors as a potential explanation for these observations. Interestingly, several of the same repeated sequences were found at multiple telomeres, often beginning at different positions within the repeat, strongly suggesting that these tandem repeats might have resulted from t-circle-mediated telomere amplification (Supp. Fig. S2D). Depending on the clone, between 12% and 22% of the total telomeric DNA was part of perfect tandem repeats, which is likely an underestimate given that we only allowed the detection of perfect matches, highlighting the importance of the inferred t-circle-mediated amplification in response to telomere uncapping.

### Telomere-uncapping-induced rearrangements are *RAD52*- and *POL32*-dependent

The Y’ rearrangement events and telomere recombination signatures we observed suggest a key role of homologous recombination (HR) pathways. To investigate the involvement of HR factors, we deleted *RAD52* and *POL32* in the *cdc13-1* strain. We exposed *cdc13-1 rad52Δ* and *cdc13-1 pol32Δ* cells to 24 h of telomere uncapping followed by recovery at 23°C. A mild ∼2-fold decrease in survival in *cdc13-1 rad52Δ* and a ∼10-fold decrease in survival in *cdc13-1 pol32Δ* compared to wild-type were observed (Supp. Fig. 3A). TRF Southern blot analysis showed that none of the *rad52Δ* and *pol32Δ* survivor clones presented either Y’-associated rearrangements or telomere elongation (Fig. 5A and B). We then performed Nanopore sequencing and genome assembly on survivor clones of both genotypes to better characterize potential rearrangements that might have escaped detection by Southern blot. The genomes of the *cdc13-1 rad52Δ* and *cdc13-1 pol32Δ* strains almost perfectly matched the control one in telomeric regions, and the rare alterations affecting Y’ elements were all Y’ losses (Fig. 5E-F). In one *pol32Δ* survivor clone, we found that chromosome III circularized using *HML* and *HMR* for homology. Circularization of chromosome III has previously been described in the context of mating-type interconversion events and recombination between mating-type loci (Klar et al., 1983; Strathern et al., 1979). Interestingly, while the *cdc13-1 rad52Δ* control strain and survivors showed modest variations in telomere length, telomeres in *cdc13-1 pol32Δ* were already longer at 23°C, consistent with a previous report (Gatbonton et al., 2006), and telomere-uncapping-induced survivors displayed highly variable telomere lengths (Fig. 5A and B, Supp. Fig. 3B and C). Overall, these results indicate that telomeric and subtelomeric rearrangements strongly depend on Rad52 and Pol32, while survival to transient telomere uncapping is partially independent of Rad52 and Pol32.

**Figure 5.**
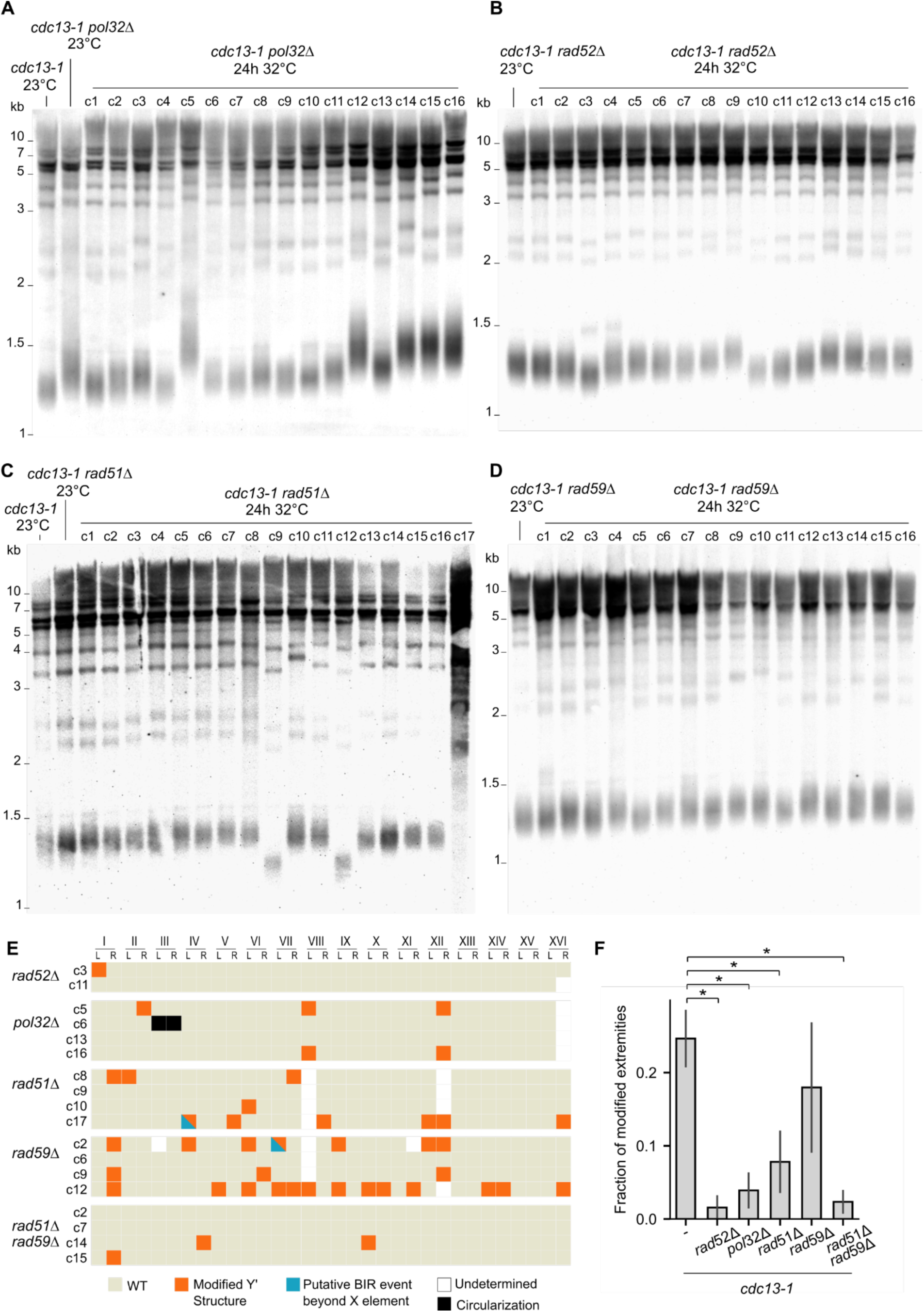
*RAD52* and *POL32* are essential for telomere-uncapping-induced rearrangements, while *RAD51* and *RAD59* are partially required. (A-D) TRF Southern blots of *cdc13-1 pol32Δ* (A), *cd13-1 rad52Δ* (B), *cdc13-1 rad51Δ* (C), and *cdc13-1 rad59Δ* (D) survivor colonies. Cultures from the same strains that were maintained at 23°C were used as controls. (E) Map of chromosome extremity alterations as in Fig. 3B, for sequenced survivor clones of the indicated genotype. A chromosome circularization event in one *pol32Δ* survivor is indicated as black squares. See Supp. Data 2 for the representation of all extremities across all sequenced clones. (F) Fractions of extremities with altered sequence or structure in the indicated strains. A statistically significant decrease was observed for the *rad52Δ*, *pol32Δ*, *rad51Δ* and *rad51Δ rad59Δ* mutants (Student’s t-test, ⁎: p-value < 0.05).

### Rad51 and Rad59 are partially involved in Y’ recombination

To further dissect the molecular pathways underlying Y’-associated and type-II-like rearrangements, we tested the roles of Rad51 and Rad59. We exposed the *cdc13-1 rad51Δ* and *cdc13-1 rad59Δ* mutants to transient telomere uncapping and observed a ∼2-fold decrease in survival in both mutants, further suggesting a protective role of homologous recombination at uncapped telomeres (Supp. Fig. S3A).

Next, we examined the structural changes in survivors using TRF Southern blotting. Only 1 out of 27 *rad51Δ* survivors and 0 out of 16 in *rad59Δ* survivors displayed a type-II-like pattern (Fig. 5C and D). While the sample size limited the statistical power, these results suggest the involvement of Rad51 and Rad59 in type-II-like formation (p-value = 0.11 and 0.08, respectively, Fischer’s exact test). Furthermore, the effects of both proteins became clearer when type-II-like signature and X band alterations were considered simultaneously: in *cdc13-1*, 18 out of 30 clones displayed alterations in TRF Southern blot, whereas only 5 out of 26 *cdc13-1 rad51Δ* clones (p-value = 0.002, Fischer’s exact test) and 4 out of 16 in *rad59Δ cdc13-1* clones (p-value = 0.03, Fischer’s exact test) were affected. Additionally, Nanopore sequencing and analysis of genome assemblies of a subset of clones confirmed a strong decrease in Yʹ recombination frequency in *rad51Δ*, whereas the effect in *rad59Δ* appeared milder (Fig. 5E and F). We then constructed the *rad51Δ rad59Δ* double mutant and found a very low frequency of modified subtelomeres, which was lower than that of each single mutant and at a level similar to that of *rad52Δ*, based on TRF Southern blot analysis and Nanopore sequencing (Fig. 5E and F, and Supp. Fig. S3F). The few remaining Y’ alterations were all Y’ losses, which did not necessarily depend on recombination. In contrast, only 2 out of 10 Y’ alterations in *rad51Δ* were losses.

Intriguingly, 2 *rad51Δ* survivors exhibited a marked reduction in telomere length (Fig. 5C and Supp. Fig. S3D). Telomere length distribution analysis of clone c9 revealed a decrease from the expected 350 bp to ∼200 bp across all chromosome ends (Supp. Fig. S3D), suggesting a potential change in global telomere length regulation in this survivor. In contrast, the telomere length distributions of the *rad59Δ* and *rad59Δ rad51Δ* survivors were not altered compared to control (Fig. 5D and Supp. Fig. S3E).

Taken together, these findings suggest that Rad51 and Rad59 both contribute to the telomeric and subtelomeric rearrangements found in Y’-associated and type-II-like survivors, and function in at least partially independent pathways.

### Type-II-like survivors are resistant to a second telomere uncapping

To test whether recombination at telomeres and subtelomeres allows yeast cells to resist telomere uncapping, we challenged 4 Y’-associated survivors and 3 type-II-like survivors with a second 24-h incubation at 32°C. While both survivor types grew normally at 23°C, only the Y’-associated survivors displayed a loss of viability comparable to that of the original *cdc13-1* control after the second stress (Fig. 6A). In contrast, all 3 type-II-like survivors maintained robust growth after 24 h at 32°C, showing little to no survival defect. We wondered whether the DNA damage checkpoint pathway in type-II-like survivors might be dysfunctional, thus supporting cell growth despite telomere uncapping. The type-II-like survivor cells arrested in G2/M after 3 h at 32°C to a similar extent as the initial *cdc13-1* strain did, as quantified by microscopy (Supp. Fig. S4), indicating DNA damage checkpoint activation in response to telomere uncapping.

**Figure 6.**
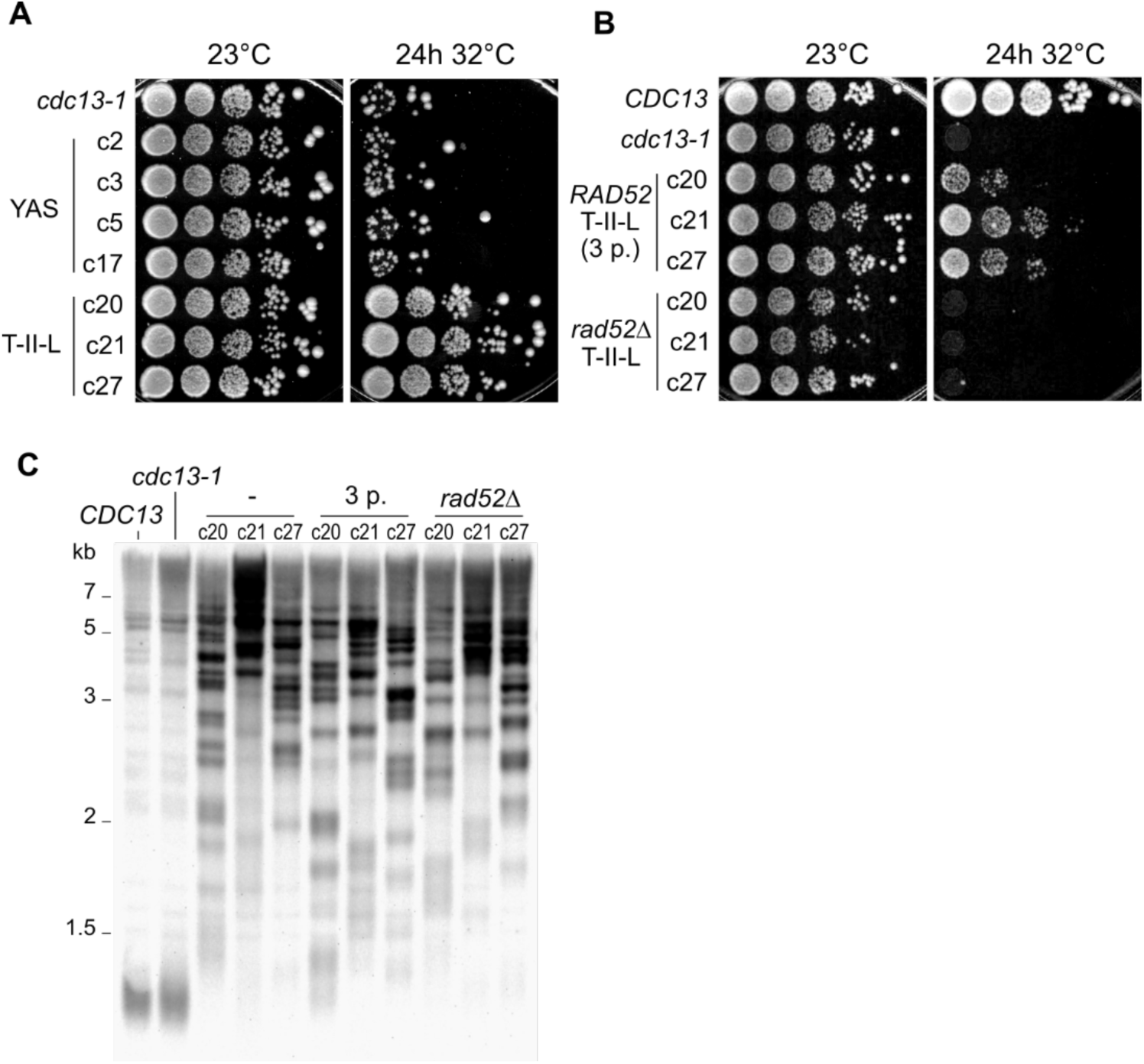
T-II-L survivors show *RAD52*-dependent ability to resist second uncapping. (A) Spot assay showing survival and growth either at a constant 23°C (left) or after a second 24-h telomere uncapping (right) for the indicated strains. (B) Same as (A) for the indicated strains. “*RAD52* T-II-L (3 p.)” c20, c21 and c27 correspond to the same strains as in (A) but were passaged 3 more times. (C) TRF Southern blots of the indicated strains. The type-II-like clones c20, c21 and c27 were analyzed once they were isolated and after 3 additional passages (“3 p.”). *RAD52* was deleted in the same clones, and the transformation and selection procedure required the equivalent of 3 passages. See Supp. Fig. S5B for reordered lanes grouped by clone.

To assess whether recombination contributed to this observed resistance, we deleted *RAD52* in the 3 previously analyzed type-II-like survivor strains. As we observed that long telomeres in type-II-like strains shortened over successive passages (Supp. Fig. S5A), we accounted for the cell divisions occurring during the transformation procedure to delete *RAD52* by passaging the control *RAD52* parents 3 times. Strikingly, the *rad52Δ* strains completely lost resistance to telomere uncapping (Fig. 6B), indicating that recombination was required for this protective effect. Importantly, this loss was not attributable to a reduction in telomere length, as TRF Southern blotting showed that the *rad52Δ* mutants retained long telomeres comparable to those of their parental strains (Fig. 6C and Supp. Fig. S5B).

Notably, the *RAD52* type-II-like strains shown in Fig. 6B exhibited reduced survival compared to those in Fig. 6A, which was particularly apparent for clone c20. Since the former were passaged 3 additional times, we attributed this decreased viability to telomere shortening that occurred during the 50–75 divisions, including potentially sporadic events of rapid telomere deletion. After 11 passages, the gradual telomere shortening led to less defined migrating bands and heterogeneity in the population of cells (Fig. S5A).

Collectively, these findings demonstrate that, together, Rad52 and long telomeres protect against a second uncapping.

## Discussion

How the genome is affected by telomere uncapping remains an outstanding question in the telomere field. Long-term survivors of permanent telomere uncapping have been previously selected after many passages and in the context of mutated checkpoint or DNA processing pathways, thus offering little information regarding the immediate molecular and cellular response to telomere uncapping in checkpoint-proficient conditions. Using a transient uncapping experimental scheme combined with molecular methods and long-read Nanopore sequencing, we provide a comprehensive characterization of the genome rearrangements following telomere uncapping, their genetic requirements and their potential functional role in protecting against further telomere deprotection.

### Early onset of genomic instability even with a functional checkpoint

Our experimental scheme allowed us to show that genome rearrangement can arise as early as during the first cell cycle after telomere uncapping, as suggested by the presence of sharp shifted PFGE bands (Fig. 2A) indicating rearrangements present in the whole clonal population. Genomic heterogeneity in *cdc13-1 cdc5-ad* survivor subclones indicates that genome instability persists for at least two cell divisions after uncapping is relieved and cells resume their cell cycle. We thus propose that transient telomere uncapping initiates a cascade of recombination events, with the first rearrangements arising within a few hours and others unfolding during the recovery phase in the progeny. Importantly, rearrangements occurred in checkpoint-proficient cells, suggesting that they did not result from uncontrolled cell cycling with broken DNA, mis-segregated chromosomes, or other abnormal structures, but rather from a canonical response to ssDNA exposed at uncapped telomeres. The use of a checkpoint-proficient background in our study stands in contrast with previous works where Cdc13-independent survivors and telomere elongation could be observed after extensive incubation at restrictive temperatures only if checkpoint or DNA processing pathways are simultaneously mutated (Grandin, 2001; Grandin & Charbonneau, 2003, 2013; Zubko & Lydall, 2006).

Therefore, transient telomere deprotection alone can be a source of genome instability, a finding with important ramifications for understanding the potential contribution of partial and transient telomere deprotection during the initial steps of tumorigenesis when checkpoints are potentially functional.

### Mechanisms underlying telomere-uncapping-driven rearrangements

Since transient-telomere-uncapping-driven telomeric rearrangements are dependent on *RAD52* and *POL32*, we propose that homologous recombination and BIR are the main molecular mechanisms underlying their formation. Additionally, both Rad51 and, to a lesser degree, Rad59 are important for Y’-associated rearrangements, especially amplifications. However, the Rad51 requirement is partial, since tandem amplification of up to 8 Y’ elements could still be observed in *cdc13-1 rad51Δ* survivors, which stands in contrast to the strictly Rad51-dependent Y’ amplification observed in telomerase-negative type I survivors (Chen et al., 2001; Teng et al., 2000). Interestingly, the fraction of type-II-like survivors measured in the *cdc13-1 rad51Δ* strain is not greater than in the *cdc13-1* strain —if anything, they tend to be less frequent in the absence of *RAD51*—, another major difference compared to telomerase-negative type II survivors, which are strongly increased in the absence of *RAD51*. We found no case of type-II-like telomere elongation in *cdc13-1 rad59Δ* survivors, suggesting that, as for telomerase-negative survivors, Rad59 might promote Rad52-mediated annealing and recombination between TG_1-3_ sequences (Churikov et al., 2014; Kockler et al., 2021), possibly through a Rad51-independent BIR pathway that relies on microhomologies (Ira & Haber, 2002). The additive effect of deleting *RAD51* and *RAD59* indicates that they act on independent pathways, at least partially.

With respect to telomere elongation in type-II-like survivors, our detailed sequence analyses strongly suggest the use of t-circles as templates for rolling circle amplification, resulting in perfect tandem repeats of specific motifs of 50-600 bp. Overall, t-circle-mediated telomere elongation accounts for ∼10-20% of all telomere sequences in type-II-like survivors and most likely more. Interestingly, t-circles also contribute to telomere elongation in telomerase-negative type II survivors (Aguilera et al., 2022; Larrivée & Wellinger, 2006), suggesting a common mechanism for generating very long telomeres regardless of the initial triggering signal.

Y’-associated survivors represent the most frequent survivor outcome of transient telomere uncapping, and we propose that extensive resection following telomere uncapping favors Y’ rearrangements owing to the homology length these elements provide and to the presence of internal telomeric sequences, as well as their organization as tandem repeats at some subtelomeres. When the homology used is within the Y’ element itself, new hybrid Y’ elements can emerge, suggesting that this process might participate in Y’ element evolution and diversification. Telomere elongation would occur less frequently and later after telomere uncapping, but not always, since we observed rearrangement patterns involving > 1 kb-long telomere sequences interspersed between Y’ elements (Fig. 3D), suggesting cases where telomere elongation precedes the invasion into an interstitial telomeric sequence and subsequent Y’ rearrangements.

Overall, extensively resected uncapped telomeres engage in multiple homology-dependent mechanisms, resulting in Y’-associated rearrangements, telomere elongation, and even more distal subtelomere duplication and translocation. While some similarities can be drawn with the mechanisms and the genetic requirements of post-senescence telomerase-independent survivors, significant differences in the response to telomere uncapping and critically short telomeres are also observed and might be due to differences in the initial length and state of the telomeres, their processing, and the signaling pathways they activate.

### Survival to transient telomere uncapping is partially dependent on homologous recombination

Loss of viability is observed when telomeres are uncapped for more than 6 h. While the exact causes of cell death remain unclear, survival partially depends on recombination factors and is associated with subtelomere and telomere rearrangements. However, altered Y’ element organization in Y’-associated survivors, the most frequent rearrangement pattern, does not confer any selective advantage in response to a second round of transient telomere uncapping. We thus speculate that recombination factors might directly promote survival by engaging molecularly with chromosome extremities, regardless of the outcome of the molecular transaction. For example, by directly interacting with ssDNA, Rad52, Rad51 and Rad59 might stabilize the end structure and limit excessive checkpoint activation, thereby acting as a protective cap for the telomeres.

On the other hand, we demonstrate that the massively elongated telomeres in type-II-like survivors strongly promote survival to a second telomere uncapping event. This finding is consistent with previous studies that used permanent telomere uncapping protocols, often together with additional mutations in checkpoint or resection pathways (Grandin, 2001; Grandin & Charbonneau, 2003; Zubko & Lydall, 2006), where survivor cells systematically displayed very long and heterogeneous telomeres. However, we show that an active *RAD52*-dependent homologous recombination pathway is required for type-II-like cells to survive a second transient telomere uncapping. A parallel can be drawn with observations that deletion of *RAD52* in type II telomerase-negative survivors very quickly leads to loss of viability, despite the presence of long telomeres(Teng & Zakian, 1999). This finding is consistent with our model, which proposes that the presence of recombination factors at telomeres or their activity is important for promoting survival to telomere uncapping. In type-II-like survivors, as compared to the parental strain, the very long telomeres would provide much more homology for Rad52-dependent annealing, thus promoting survival. Conceptually, we thus propose that recombination factors or the molecular structures formed by long telomeres undergoing recombination serve as an alternative cap for telomeres when Cdc13 is dysfunctional.

## Material and Methods

### Yeast strains

All strains are from the W303 background (*ura3-1 trp1-1 leu23,112 his3-11,15 can1-100*) corrected for *RAD5* and *ADE2* (Supp. Table 1). Most strains carry the *cdc13-1* allele and were grown routinely at the permissive temperature of 23°C in YPD (yeast extract, peptone, dextrose) media. Deletion strains were created using PCR-based methods as described in (Longtine et al., 1998). Point mutations were introduced using Cas9-mediated gene targeting as described in (Anand et al., 2017).

### Transient uncapping survival assay

Cells were inoculated in YPD and grown overnight at 23°C. The culture was then diluted and grown in exponential phase until it reached a concentration of 1-2 x 10^7^ cells/mL. Between 5 x 10^5^ and 5 x 10^6^ cells were plated on solid YPD media preheated at 32°C and then kept at 32°C for 24 h before the temperature was shifted back to 23°C until colony formation (∼3 days). Another plate was inoculated with ∼500 cells of the same liquid culture and kept at 23°C to measure plating efficiency and to be used for normalization. The number of cells plated was dependent on the expected survival frequency of each strain and was chosen in order to yield between 50 and 300 individual colonies. The colonies on the plate that was placed transiently at 32°C and on the plate kept at 23°C were then counted to calculate the survival frequency. Colonies randomly selected for further investigation were subsequently subcloned and then grown at 23°C.

### Genomic DNA extraction

Two genomic DNA extraction protocols were used in this study for subsequent Nanopore sequencing. Details of the extraction protocol used in each strain can be found in Supp. Table 2. For the first protocol, QIAGEN Genomic-tip 100/G was used. A total of 5 x 10^9^ cells from an overnight liquid culture were pelleted, washed with water, and then resuspended in 4 mL of Y1 buffer with 400 U of lyticase for 1 h at 30°C. The resulting spheroblasts were pelleted and resuspended in 5 mL of G2 buffer with 1 mg of RNase A and 1 mg of proteinase K for 1 h at 50°C. The lysate was centrifuged at 12000 g for 15 min at 4°C, and the supernatant was then passed through a Qiagen column according to the manufacturer’s instructions. For the second protocol, 1-2 x 10^9^ cells from an overnight liquid culture were pelleted, washed with 500 µL of spheroblasting solution (1 M sorbitol, 50 mM KPO_4_, 10 mM EDTA), resuspended in 500 µL spheroblasting solution supplemented with 5 µL of β-mercaptoethanol and 1.25 mg of zymolyase 20T and incubated for 30 min at 37°C. Spheroblasts were pelleted at 2500 g for 3 min, resuspended in 500 µL of lysis solution (0.1 M Tris HCL at pH 8.0, 50 mM EDTA, 0.5 M NaCl, 1% PVP40, 2.5% SDS) with 1.5 mg of RNase A, and then incubated for 45 min at 50°C. 1 mL of TE and 500 µL of 5 M potassium acetate were added and gently mixed in before 2 centrifugation steps of 10 min at 9500 g at 4°C to clarify the supernatant. DNA was precipitated from the supernatant with an equal volume of isopropanol and washed with 70% cold ethanol before final elution in 40 µL of H_2_0.

### Pulsed field gel electrophoresis

Agarose plugs containing 2 x 10^8^ yeast cells were prepared as described in (Török et al., 1993) and sealed in a 0.5X TBE (0.89 M Tris, 0.89 M boric acid, and 20 mM EDTA) 1% Seakem GTC agarose gel. PFGE was run on the CHEF-DRII system (Bio-Rad) with the following program: 6 V/cm for 10 h with a switching time of 60 sec followed by 6 V/cm for 17 h with a switching time of 90 sec. The reorientation angle was set to 120° throughout the entire run.

### Library preparation and sequencing

The sequencing libraries were prepared following Oxford Nanopore Technologies (ONT) protocols for genomic DNA without preamplification and using LSK109 or LSK110 kits. Genomic DNA was enriched for long fragments using the SRE kit (PacBio). DNA libraries were sequenced on R9.4 or R10.4 Nanopore flow cells in either a MinION Mk1C or PromethION P2 sequencer, respectively, with default parameters on the MinKNOW operating software. For each run, 500 ng of DNA was loaded and sequenced for 8–24 h. In case of DNA reloading, the flow cell was washed using ONT’s wash kit following the instructions.

### TRF Southern blot

Genomic DNA was extracted from cultures using a standard phenol:chloroform:isoamyl (25:24:1) purification procedure and isopropanol precipitation. A sample of 2 µg of genomic DNA was digested with XhoI, and the products were ethanol-precipitated, resuspended in loading buffer (purple gel loading dye 6X, New England Biolabs), and resolved on a 1.1% agarose gel for 6 h at 3.6 V/cm. The gel was then soaked in a denaturation bath (0.4 M NaOH, 1 M NaCl) for 20 min and transferred by capillary action to a charged nylon membrane (Hybond XL, GE Healthcare). A telomere-specific oligonucleotide probe (5ʹ-CCCACCACACACACCCACACCC-3ʹ) biotinylated at both ends was used for hybridization. After hybridization of the probe, the membrane was washed 3×5 min in wash buffer (58 mM Na2HPO_4_, 17 mM NaH_2_PO_4_, 68 mM NaCl, 0.1% SDS). The membrane was next processed for detection with 3 successive incubations (5, 5 and 30 min) in blocking buffer (Thermo Scientific, Nucleic Acid Detection Blocking Buffer) before a 30 min incubation with alkaline phosphatase-conjugated streptavidin (Invitrogen) diluted in blocking buffer (0.4 g/mL). The membrane was then washed again 3 × 5 min in wash buffer, incubated for 2 × 2 min in assay buffer (0.1 M Tris, 0.1 M NaCl pH 9.5) and for 5 min in CDP-Star substrate (Applied Biosystems) before being imaged with a Bio-Rad ChemiDoc Touch device by chemiluminescence.

### Whole genome assemblies

Data from sequencing was basecalled using Guppy with model dna_r9.4.1_450bps_sup.cfg for R9 data and Dorado for with model dna_r10.4.1_e8.2_400bps_sup@v5.0.0 for R10 data. Reads were filtered using Filtlong to keep only the best 500-700 Mb using the filtering parameters --length_weight 6 -- mean_q_weight 3 --window_q_weight 1 –min_length 1000. The genomes were assembled using Canu 2.2 and Flye 2.9.1, using standard parameters for Canu and the option --nano-hq for Flye, then polished using one round of Racon v1.5.0 followed by 2 rounds of Medaka (v1.11.3 for R9 data, v2.0.1 for R10 data). The output assemblies were mapped against the S288C reference genome for contig name annotation, and short contigs (<70 kb) and contigs corresponding to mitochondrial DNA were removed. To obtain the final assembly, reads were mapped against Canu and Flye assemblies, telomeric regions were manually inspected using Tablet, and the best contigs were retained for each chromosome. Trimming of contigs was manually performed when overassembly at telomeres was detected through loss of coverage. The statistics of each sample for the reads and assemblies can be found in Supp. Table 2.

### Annotation of telomeric regions

Telomeres were detected on assemblies using Telofinder (O’Donnell 2023) (https://github.com/GillesFischerSorbonne/telofinder). X elements were detected in the assemblies using LRSDAY 1.7.2 (Yue & Liti, 2018). Y’ elements were detected by aligning known Y’ elements from the reference genome against the considered assembly using BLAST 2.12.0, and then taking the union of all alignments whose lengths were greater than 200 bp.

### Y’ element labelling and clustering

Because Nanopore sequencing makes most errors in homopolymers, the detected Y’ sequences were transformed by condensing all homopolymers of length >4 into homopolymers of length 4. Doing so increased robustness and yielded a 100% similarity between the Y’ elements of the 2 independent assemblies of our control strain. The similarity measure was derived from BLAST alignment results, using the best alignment between each pair of sequences. The formula used was sim = %identity x alignment_length/alignement_span where alignment_span is the length of the alignment plus the length of the unaligned flanking regions, and %identity was obtained from BLAST.

To assess the precision of our method, another set of reads was split and assembled into 2 independent assemblies, which yielded similarity values between homologous Y’ between 99.985% and 100%. Labels were then assigned to Y’ elements of our control strain. Two or more Y’ elements were given the same label if their similarity score was higher than 99.9%.

Hierarchical clustering of the Y’ elements of our control strain was performed on their similarity matrix using Seaborn’s clustermap function with default parameters. The optimal number of 11 clusters was determined by determining the elbow point of the silhouette score.

### Subtelomeric alteration detection

To detect Y’ element structure alterations, we compared the labels assigned to each element to the labels of the elements in the corresponding control strain. Any label that was present at a position in the control and not in a survivor was considered a loss. Any label present at a position in a survivor and not in the control was considered a gain.

To detect subtelomeric events not including Y’ elements or telomeres, sequences upstream of the X element at each extremity were aligned over 15 kb using BLAST against their control counterparts, and any deviation from perfect alignment was flagged for manual investigation.

To check whether a new Y’ sequence not present in the control genome could be explained by a mosaic of elements in the control genome, we used a custom python script to look for perfect alignments of size >50 between a new Y’ element and all known Y’ elements. A Y’ element was considered mosaic if it could be covered by the union of such alignments.

## Data availability

All new sequencing data (fast5, fastq and fasta files) were deposited in the European Nucleotide Archive (https://www.ebi.ac.uk/ena/ browser/home) with the project accession PRJEB93986.

## Acknowledgement

We thank Stéphane Delmas and Teresa Teixeira for their critical reviews of the paper. We thank Nicolas Agier for his help and advice with Nanopore sequencing and subsequent analyses. Work in ZX’s lab was supported by Ville de Paris (Programme Émergence(s)), the Emergence grant of Sorbonne Université, Ligue Contre le Cancer (Subvention Recherche Scientifique 2022), Fondation ARC pour la recherche sur le cancer (ARCPJA202160003865 and ARCPGA2023110007341_7967), and by Agence Nationale de la Recherche (ANR-24-CE12-7740-01 and ANR-24-CE12-0083-03).

## Author contributions

LD: Investigation, Methodology, Formal Analysis, Writing – Original Draft. CG: Methodology, Formal Analysis. OI: Investigation. JSB: Conceptualization, Supervision. ZX: Conceptualization, Supervision, Writing – Original Draft. All authors – Writing – Review & Editing.

