## Supplementary Information for "Transient telomere uncapping triggers telomeric and subtelomeric rearrangements"

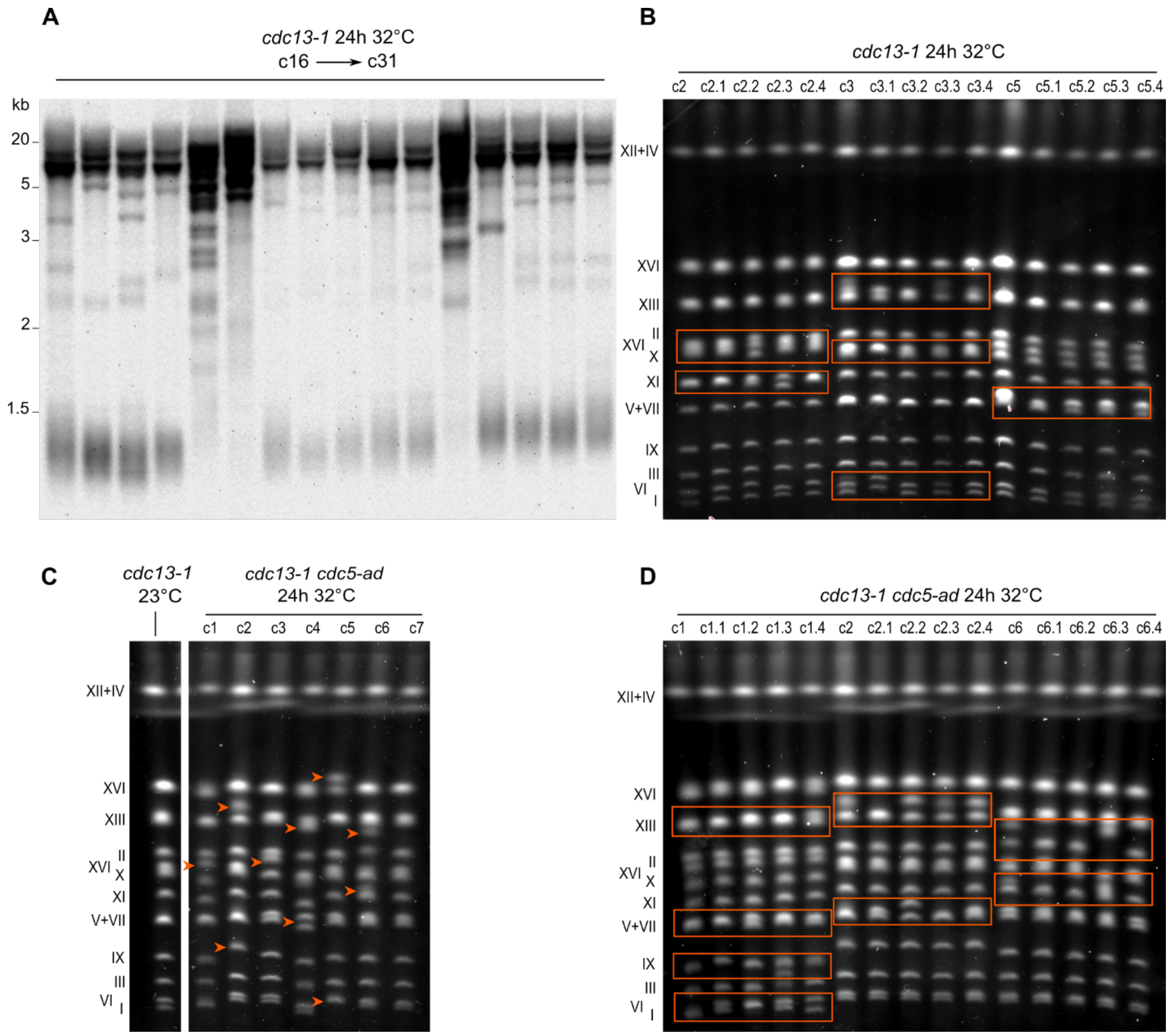


**Supplementary Figure S1. Heterogeneous genome rearrangements at telomeres and subtelomeres.**

(A) TRF Southern blot analysis of 16 *cdc13-1* survivor clones in addition to those shown in Fig. 2B.

(B) PFGE of 4 subclones of clones c2, c3 and c5. The orange boxes indicate bands with distinct migration patterns in the 4 subclones.

(C) PFGE of 7 *cdc13-1 cdc5-ad* transient uncapping survivor clones as well as a *cdc13-1* control strain grown constantly at 23°C. Compared to the control strain, 6 out of 7 survivor clones exhibited apparent chromosome size shifts, marked by orange arrows.

(D) Same as (B) in a *cdc13-1 cdc5-ad* strain.

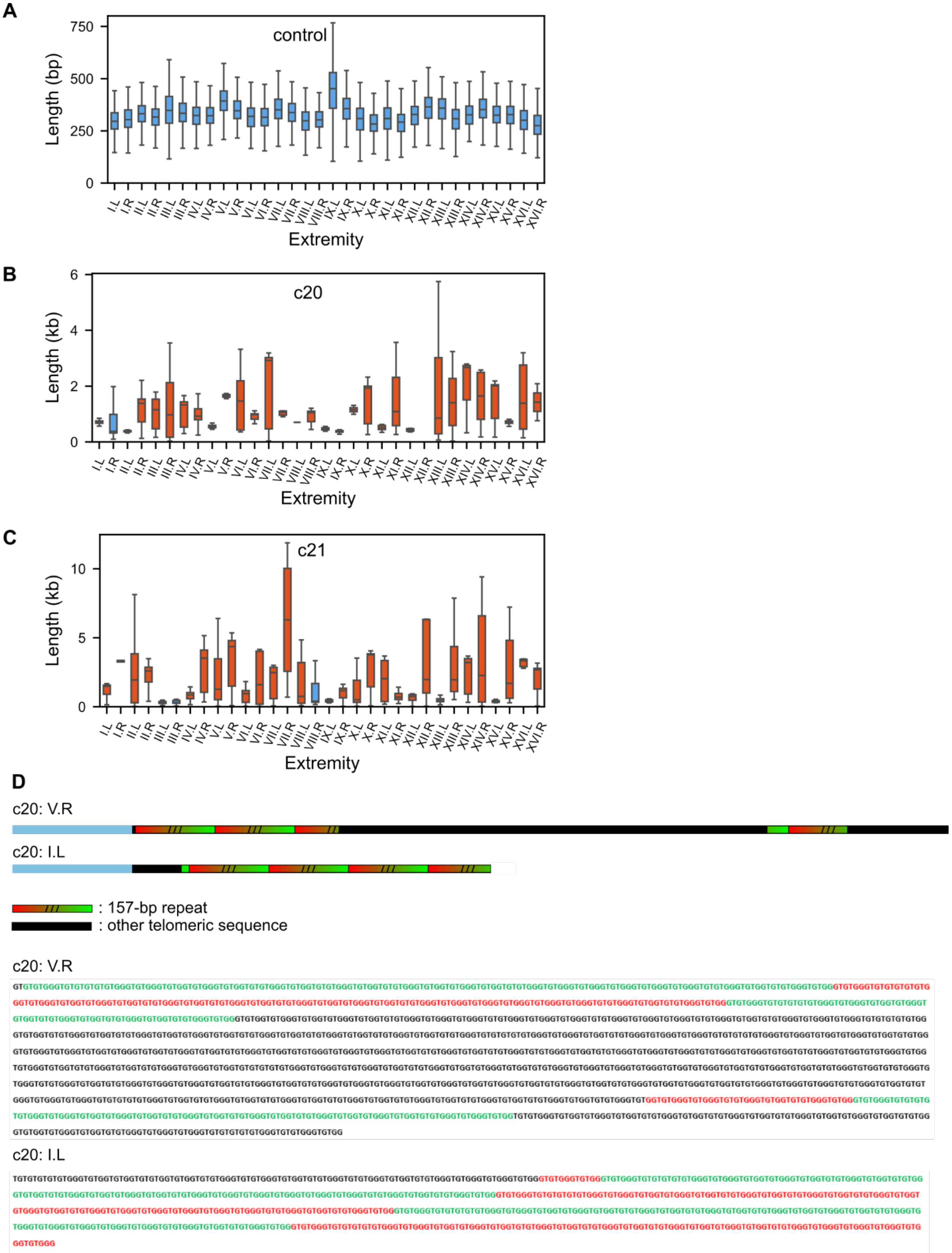


**Supplementary Figure S2. Telomeres are elongated heterogeneously in T-II-L survivors and contain large**

**perfect tandem repeats.**

(A) Boxplots representing the telomere length distribution at each chromosome extremity of the control *cdc13-1* strain grown at 23°C. Same data as in Fig. 4C left with a different scale on the y-axis.

(B-C) Same as (A) for type-II-like survivor clones c20 and c21. The blue boxplots correspond to distributions with a mean telomere length < 500 bp.

(D) Schematic representation of arrays of perfect tandem telomere repeats in type-II-like survivor clone c20. The same 157-bp repeat is found in arrays at 2 telomeres (V.R and I.L), starting at different positions in the motif. The exact sequences are shown below.

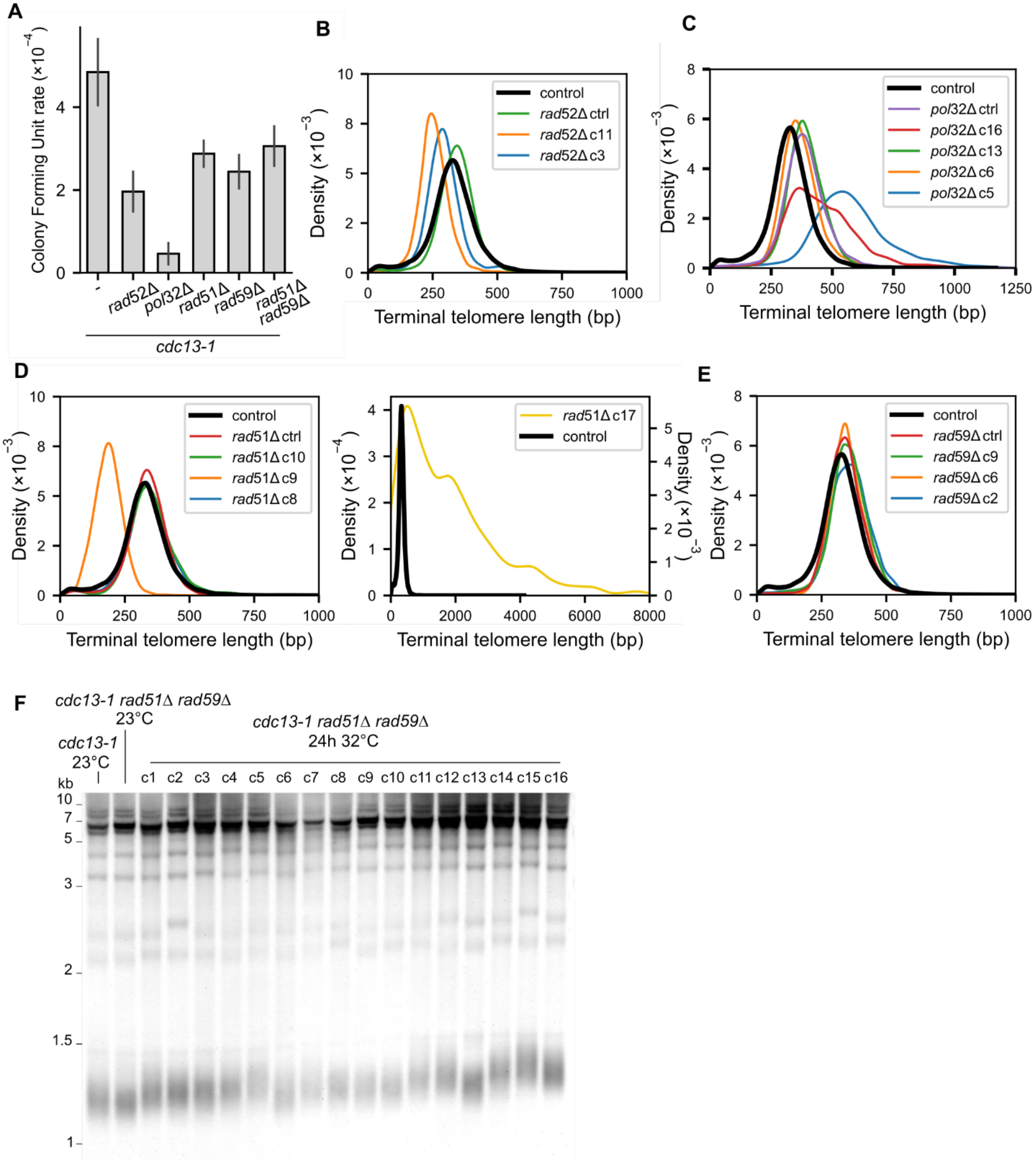


**Supplementary Figure S3. Telomere length distributions of transient uncapping survivors in the *rad52Δ*, *pol32Δ*, *rad51Δ*, and *rad59Δ* mutants.**

(A) Colony formation frequency after transient telomere uncapping and return to 23°C in the indicated mutants. A control plate with the *cdc13-1* strain was kept only at 23°C for normalization. The error bars correspond to the standard error of the mean.

(B-E) Telomere length distributions of the indicated strains derived from Nanopore sequencing reads, compared to the control (black line). In (D), the left plot displays the telomere length distribution of *rad51Δ* YAS survivors while the right plot displays the telomere length distribution of for the *rad51Δ* type-II-like survivor (c17), with a different scale for the x-axis.

A

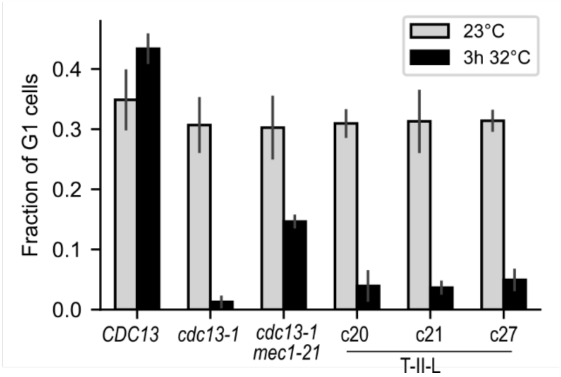


**Supplementary Figure S4. Type-II-like survivors arrest in G2/M in response to telomere uncapping.**

Fraction of unbudded G1 cells in exponentially growing cultures at 23°C or after 3 hours at 32°C. G1 cells are more abundant in the partially checkpoint-deficient mutant *mec1-21 cdc13-1* than in the control *cdc13-1*. N = 3 independent cultures for each condition.

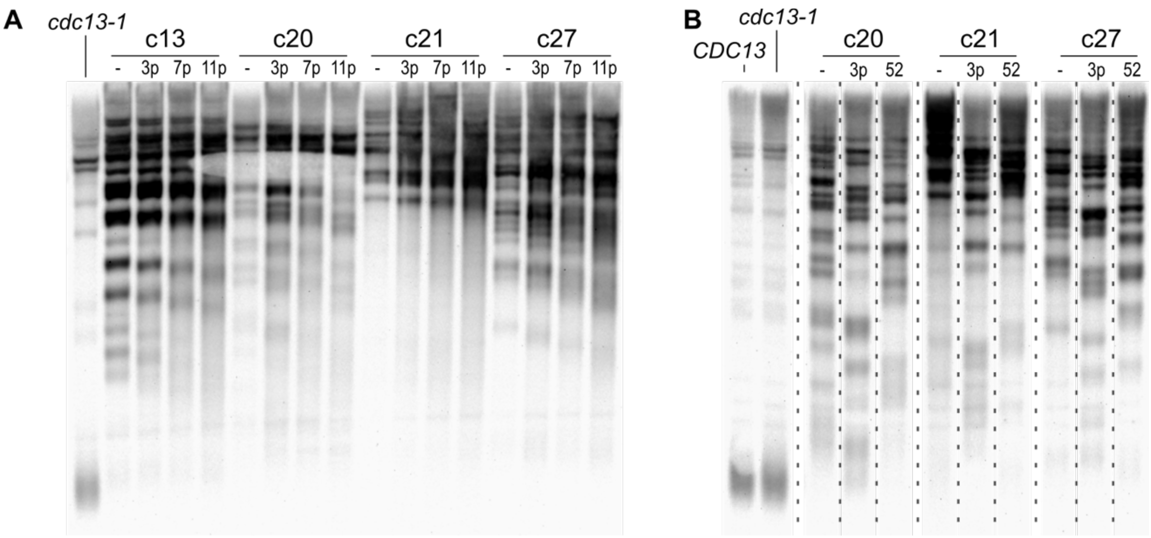


**Supplementary Figure S5. Telomere shortening of type-II-like survivors over passages.**

(A) TRF Southern blot of 4 type-II-like clones, passaged for an additional 3, 7 and 11 passages after initial subcloning. Contrary to Fig. 6C, restreaks were performed on bulk patches of cells instead of colonies.

(B) Figure identical to Fig. 6C, except that the lanes have been digitally reordered to facilitate visual interpretation per clone. “52” indicates the *rad52Δ* derivative of the indicated strain.

| **Strain** | Genotype |
|---|---|
| **yT1291** | MAT**α** *ura3-1 trp1-1 leu2-3,112 his3-11,15 can1-100* ADE2 RAD5 *cdc5::CDC5-3HA-TRP1 cdc13-1* |
| **yT1292** | MAT**α** *ura3-1 trp1-1 leu2-3,112 his3-11,15 can1-100* ADE2 RAD5 *cdc5::cdc5-ad-3HA-TRP1 cdc13-1* |
| **yZX409** | yT1291 *rad51::HIS3* |
| **yZX419** | yT1291 *rad52::HIS3* |
| **yZX424** | yT1291 *rad59::HIS3* |
| **yZX447** | yT1291 *pol32::HIS3* |
| **yZX500** | yT1291 *mec1-21* |
| **yZX503** | yT1291 *rad59::HIS3 rad51::LEU2* |

**Supplementary Table 1. Yeast strains used in this study.**

1

| Strain | Chemistry | Bases kept | Read count | median depth | N50 | PHRED quality | Nb of contigs | Asssembly size (b) |
|---|---|---|---|---|---|---|---|---|
| **yT1291** | SQK-LSK109 | 600M | 23500 | 38 | 27354 | 15.46 | 16 | 12254979 |
| **yT1291_2** | SQK-LSK109 | 700M | 10054 | 51 | 72413 | 16.39 | 16 | 12374096 |
| yT1291_survivor_c2 | SQK-LSK109 | 600M | 22814 | 35 | 27842 | 14.05 | 17 | 12381501 |
| yT1291_survivor_c3.1 | SQK-LSK109 | 600M | 23728 | 34 | 26273 | 14.65 | 17 | 12385410 |
| yT1291_survivor_c3.2 | SQK-LSK109 | 600M | 22788 | 34 | 21138 | 14.64 | 17 | 12369144 |
| yT1291_survivor_c3.3 | SQK-LSK109 | 600M | 29840 | 29 | 21138 | 14.15 | 16 | 12399802 |
| yT1291_survivor_c3.4 | SQK-LSK109 | 600M | 36991 | 29 | 16703 | 14.38 | 20 | 10968454 |
| yT1291_survivor_c5 | SQK-LSK109 | 600M | 23127 | 37 | 27912 | 14.61 | 17 | 12310970 |
| yT1291_survivor_c17 | SQK-LSK109 | 600M | 22085 | 37 | 28315 | 14.84 | 17 | 12432499 |
| yT1291_survivor_c20 | SQK-LSK109 | 581M | 42120 | 32 | 20924 | 13.38 | 17 | 12305791 |
| yT1291_survivor_c21 | SQK-LSK109 | 600M | 30013 | 31 | 22007 | 14.18 | 16 | 12368767 |

| | | | | | | | | |
|---|---|---|---|---|---|---|---|---|
| yT1291_survivor_c26 | SQK-LSK110 | 600M | 13423 | 39 | 43254 | 21.03 | 19 | 12492133 |
| yT1291_survivor_c27 | SQK-LSK109 | 600M | 39491 | 31 | 20928 | 13.37 | 17 | 12354973 |
| yT1291_survivor_c29 | SQK-LSK110 | 600M | 6324 | 42 | 93349 | 24.6 | 18 | 12759246 |
| **yZX424** | SQK-LSK110 | 600M | 9428 | 37 | 64933 | 23.8 | 17 | 12436295 |
| yZX424_survivor_c2 | SQK-LSK110 | 600M | 10639 | 27 | 54440 | 22.7 | 20 | 11167879 |
| yZX424_survivor_c6 | SQK-LSK110 | 600M | 13517 | 37 | 42869 | 24.58 | 16 | 12292235 |
| yZX424_survivor_c9 | SQK-LSK110 | 600M | 9748 | 36 | 59303 | 22.84 | 17 | 12297725 |
| yZX424_survivor_c12 | SQK-LSK110 | 600M | 11500 | 39 | 55698 | 22.62 | 17 | 12412417 |
| **yZX409** | SQK-LSK110 | 600M | 15797 | 32 | 36955 | 24.08 | 18 | 12492910 |
| yZX409_survivor_c8 | SQK-LSK110 | 600M | 11161 | 40 | 52275 | 23.75 | 17 | 12483071 |
| yZX409_survivor_c9 | SQK-LSK110 | 600M | 14151 | 34 | 40646 | 23.25 | 18 | 12669384 |
| yZX409_survivor_c10 | SQK-LSK110 | 600M | 14310 | 32 | 40646 | 23.25 | 17 | 12379387 |
| yZX409_survivor_c17 | SQK-LSK110 | 600M | 13810 | 34 | 41877 | 24.1 | 17 | 12473960 |
| **yZX419** | SQK-LSK110 | 600M | 13530 | 27 | 42948 | 20.34 | 18 | 12257221 |

| | | | | | | | | |
|---|---|---|---|---|---|---|---|---|
| **yZX419_survivor_c3** | SQK-LSK110 | 600M | 11990 | 33 | 47334 | 23.09 | 19 | 12373752 |
| **yZX419_survivor_c11** | SQK-LSK110 | 600M | 9137 | 40 | 62967 | 20.82 | 17 | 12247306 |
| **yZX447** | SQK-LSK110 | 600M | 8909 | 40 | 64933 | 23.8 | 17 | 12248378 |
| **yZX447_survivor_c5** | SQK-LSK110 | 600M | 9901 | 42 | 57975 | 24.1 | 16 | 12321768 |
| **yZX447_survivor_c6** | SQK-LSK110 | 600M | 7967 | 42 | 71286 | 24.71 | 16 | 12330222 |
| **yZX447_survivor_c13** | SQK-LSK110 | 600M | 10467 | 42 | 55385 | 23.97 | 17 | 12351550 |
| **yZX447_survivor_c16** | SQK-LSK110 | 600M | 12623 | 39 | 46147 | 20.79 | 17 | 12285666 |
| **yZX503** | SQK-LSK110 | 600M | 10706 | 39 | 55698 | 22.91 | 17 | 12535522 |
| **yZX503_survivor_c2** | SQK-LSK110 | 600M | 9242 | 38 | 64338 | 23.11 | 17 | 12359884 |
| **yZX503_survivor_c7** | SQK-LSK110 | 600M | 11479 | 38 | 53107 | 22.63 | 18 | 12562493 |
| **yZX503_survivor_c14** | SQK-LSK110 | 600M | 8335 | 40 | 70524 | 23.53 | 17 | 12425479 |
| **yZX503_survivor_c15** | SQK-LSK110 | 600M | 8177 | 40 | 71656 | 23.38 | 17 | 12382989 |

**Supplementary Table 2. Sequencing and assembly statistics.**

"Chemistry" indicates the type of flowcell and reactants used. "Bases kept" indicates the number of bases used for assembly after filtering. "Read count" indicates the number of reads used for assembly after filtering. "Median depth" indicates the median depth of filtered reads mapped to the final assembly. "N50" corresponds to the N50 of the reads used for assembly. "PHREd quality" is the average PHRED quality score of the reads used for assembly. "Nb of contigs" is the final number of contigs in assemblies, excluding mitochondrial contigs.

**Supplementary Data 1. List of all Y' elements in the cdc13-1 control strain.**

Fasta file containing the sequences of all 34 identified Y' elements of the *cdc13-1* control strain grown only at 23°C.

**Supplementary Data 2. Representations of the telomeric and subtelomeric regions of all sequenced strains and survivor clones.**

The schematic representations are aligned on the X element for each chromosome extremity. One folder corresponds to one genotype. The legend and clone IDs are shown in the image of Chr_I.left for each genotype and are applicable to extremities of the same genotype.
