## Supplementary figures and images for "Transient telomere uncapping triggers telomeric and subtelomeric rearrangements"

### Chr_I.left.png

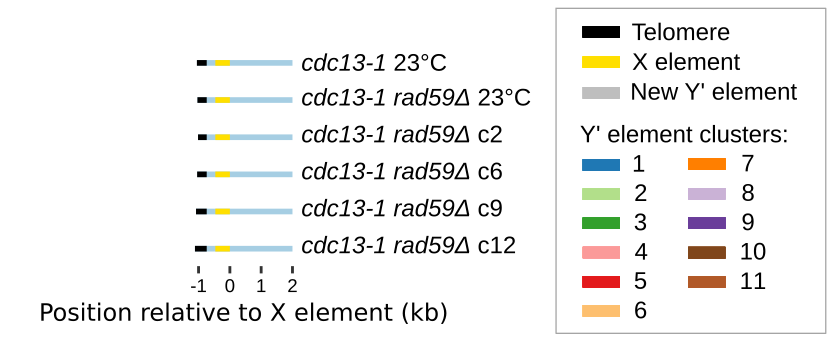

### Chr_I.left.png

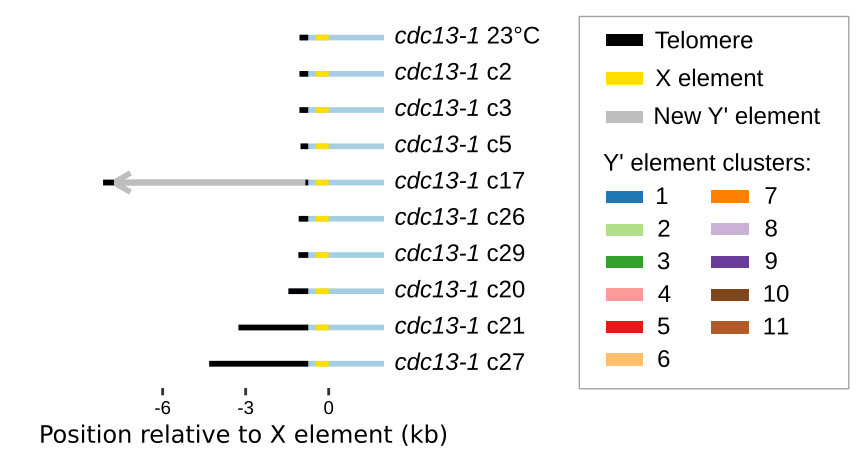

### Chr_I.right.png

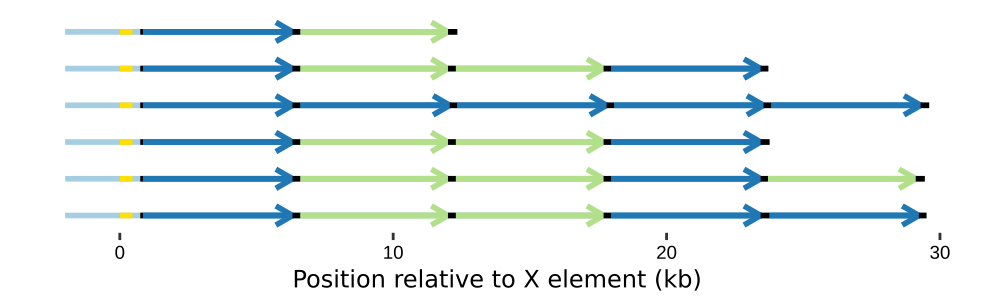

### Chr_I.right.png

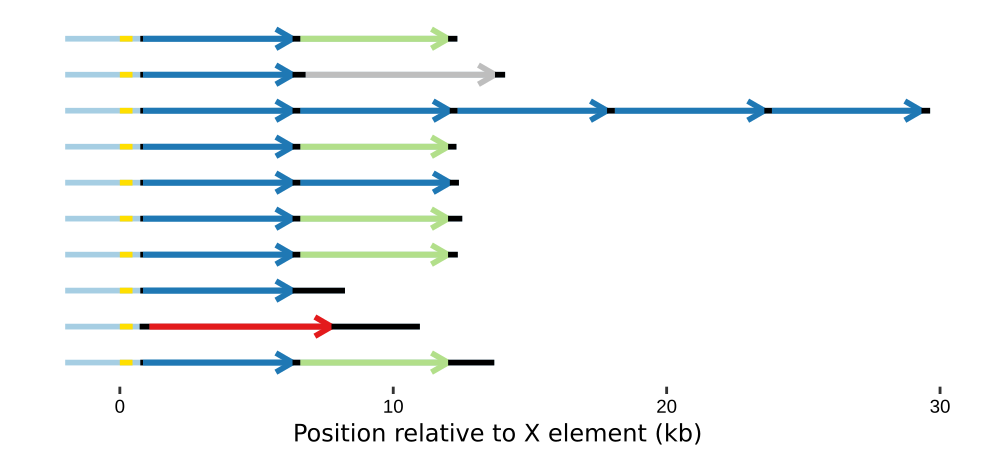

### Chr_II.left.png

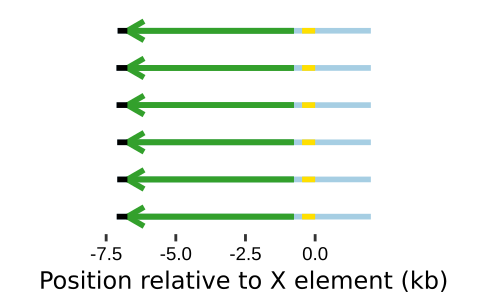

### Chr_II.left.png

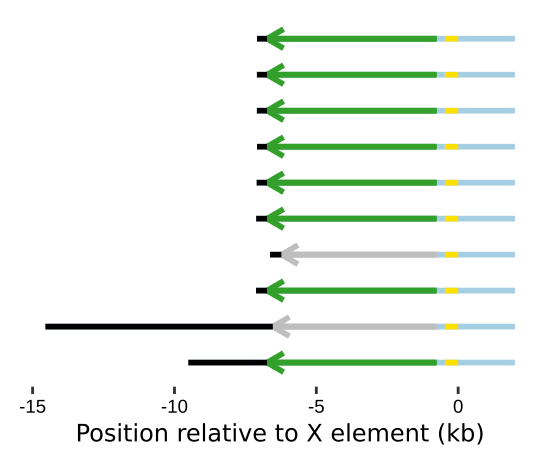

### Chr_II.right.png

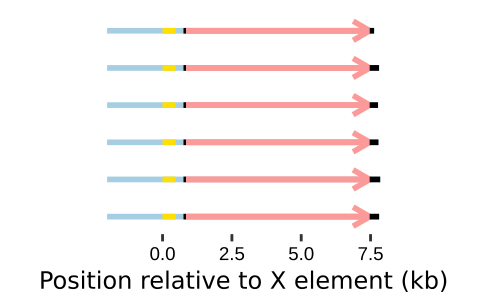

### Chr_II.right.png

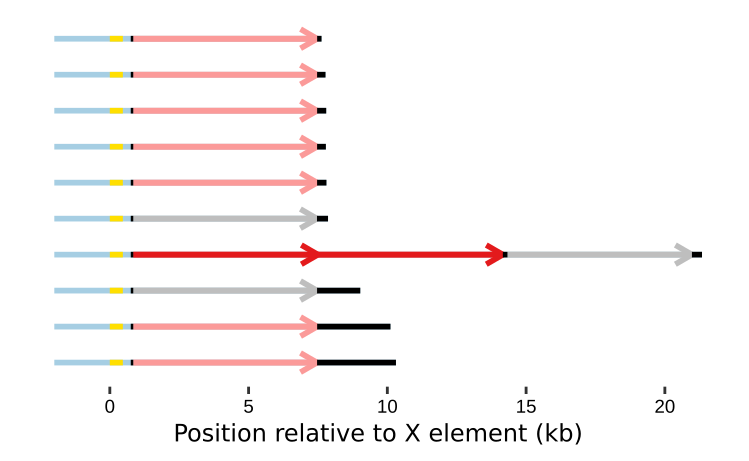

### Chr_III.left.png

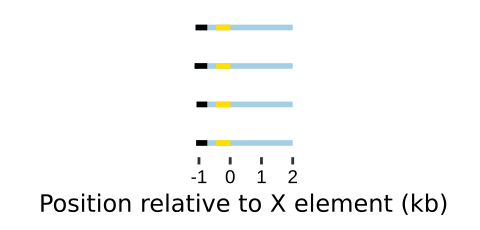

### Chr_III.left.png

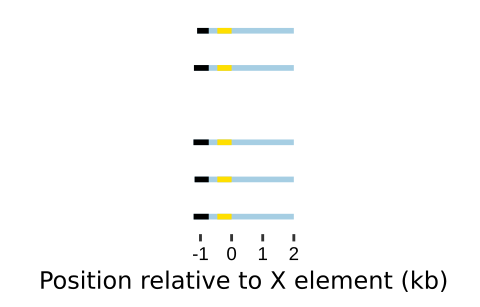

### Chr_III.left.png

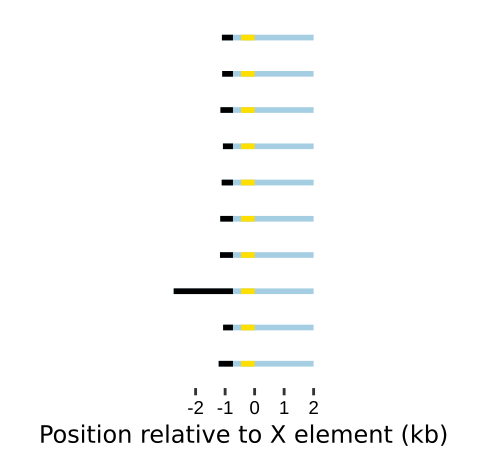

### Chr_III.right.png

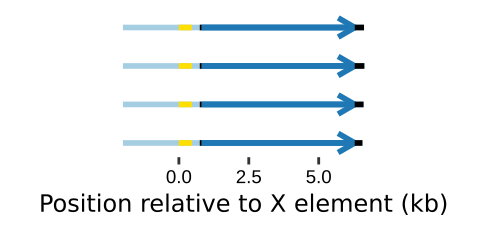

### Chr_III.right.png

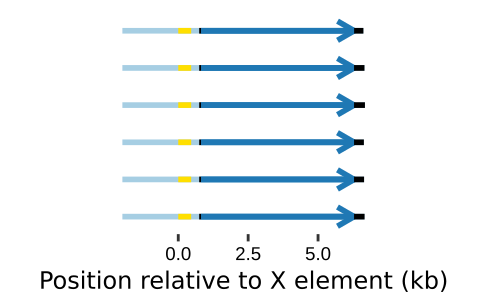

### Chr_III.right.png

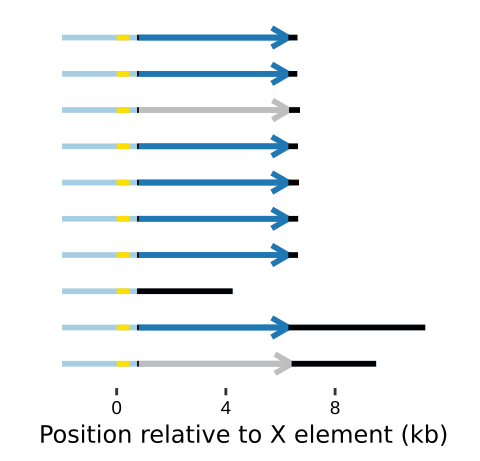

### Chr_IV.left.png

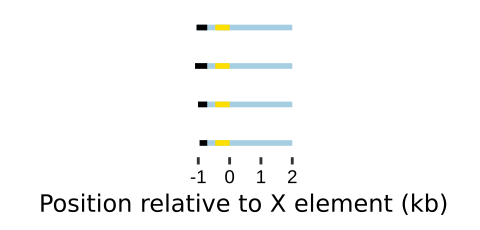

### Chr_IV.left.png

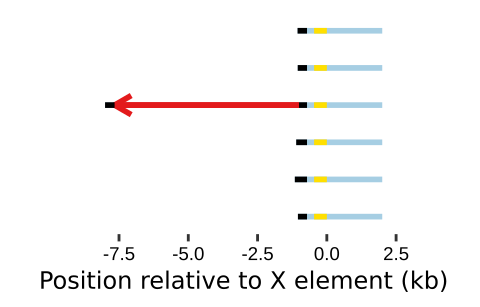

### Chr_IV.left.png

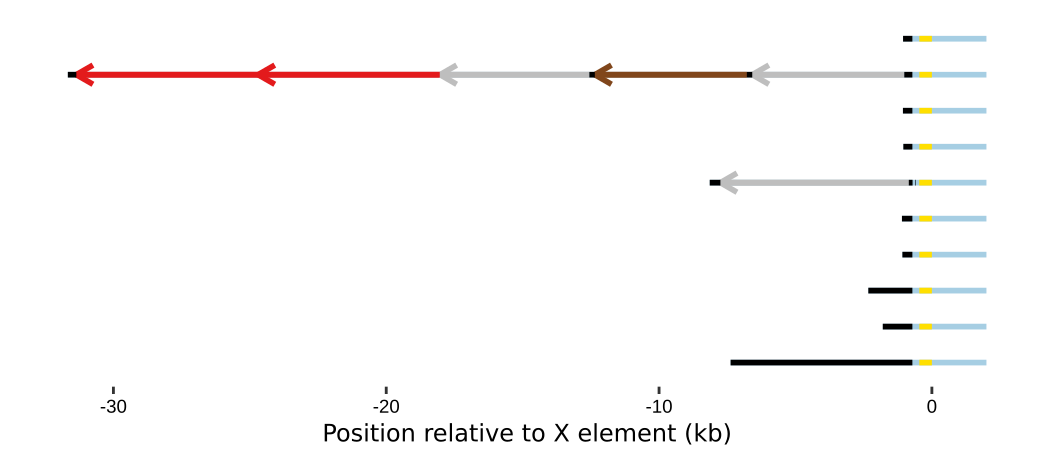

### Chr_IV.right.png

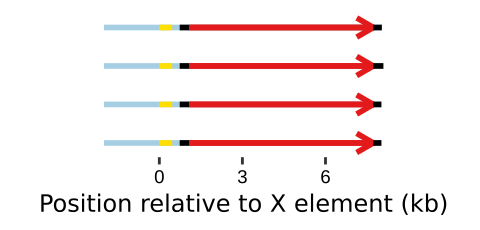

### Chr_IV.right.png

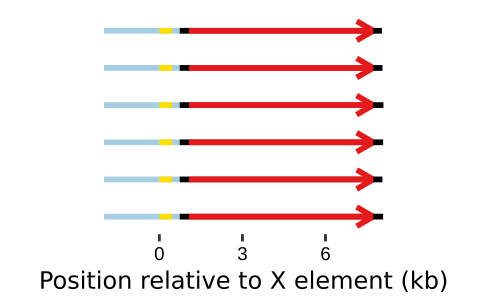

### Chr_IV.right.png

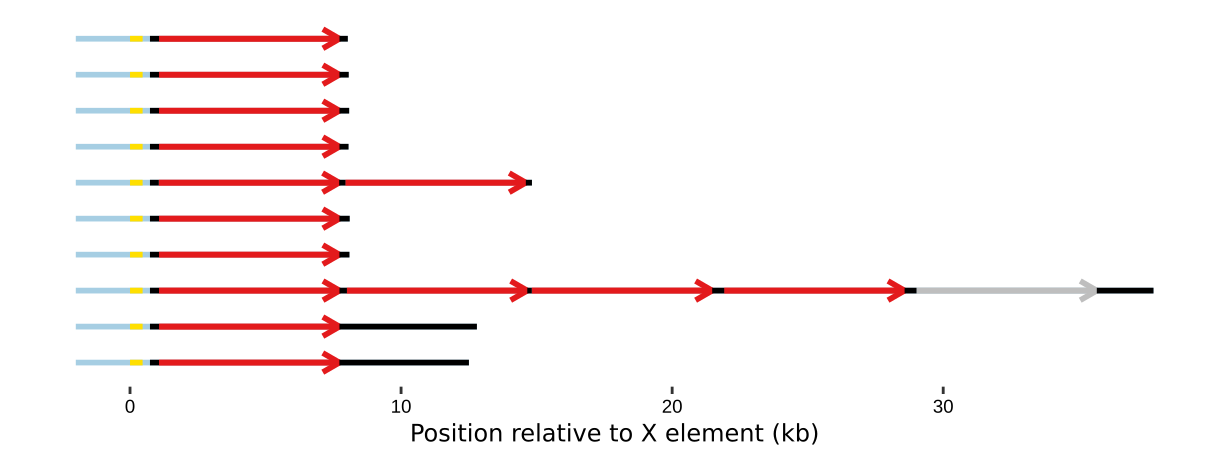

### Chr_IX.left.png

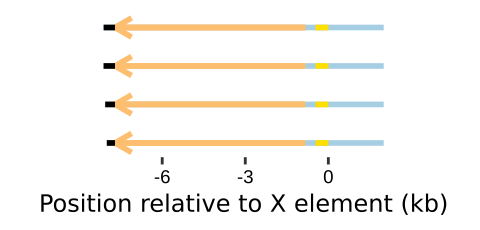

### Chr_IX.left.png

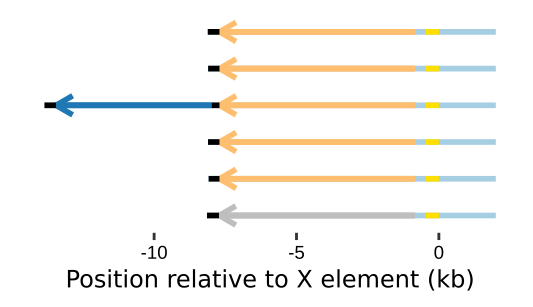

### Chr_IX.left.png

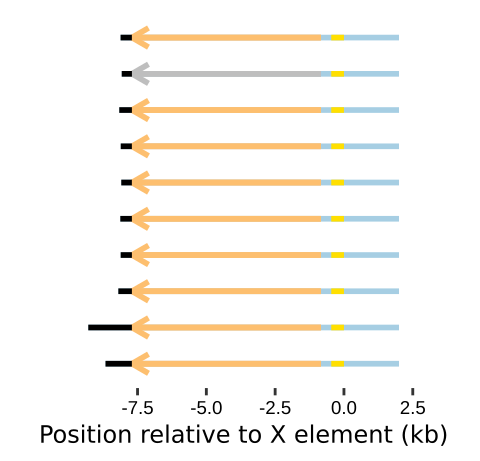

### Chr_IX.right.png

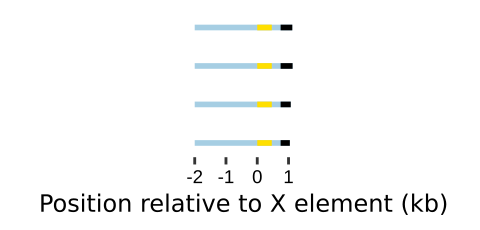

### Chr_IX.right.png

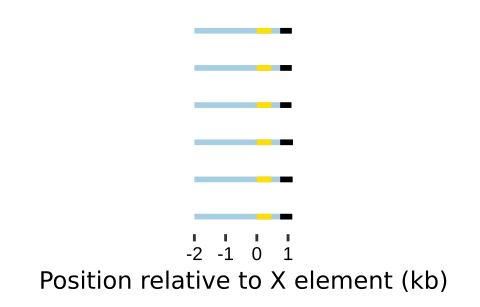

### Chr_IX.right.png

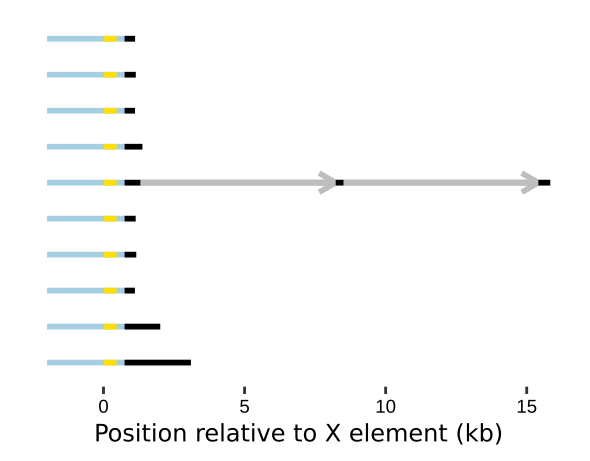

### Chr_V.left.png

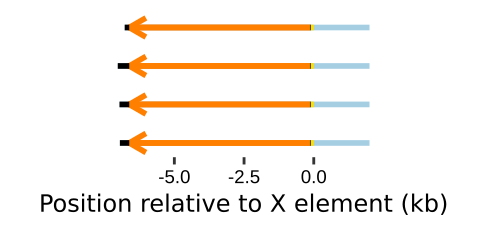

### Chr_V.left.png

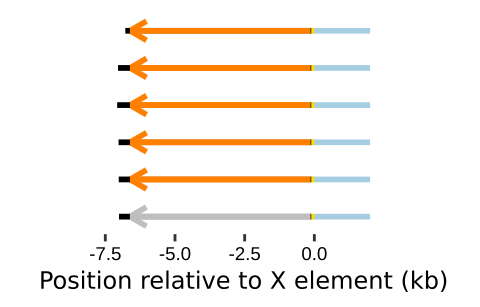

### Chr_V.left.png

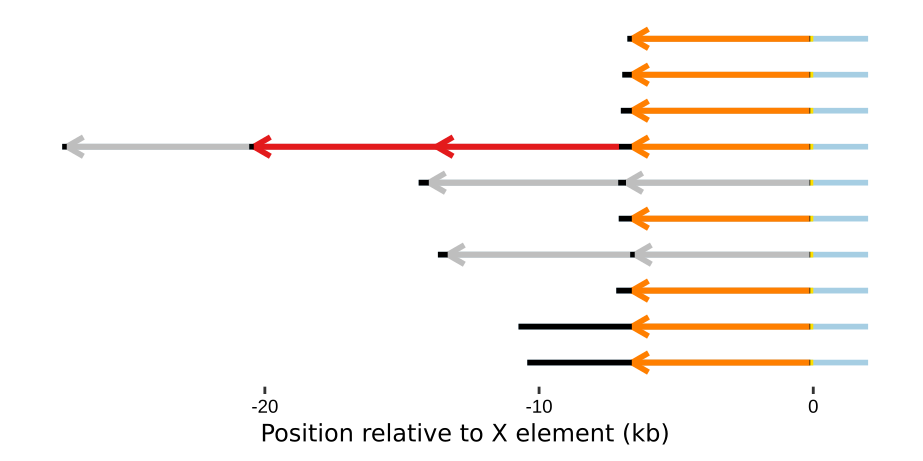

### Chr_V.right.png

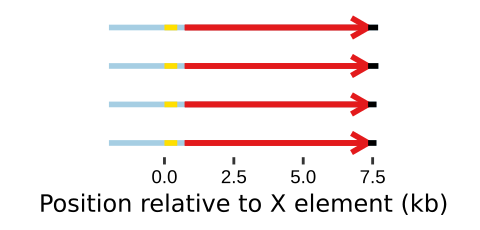

### Chr_V.right.png

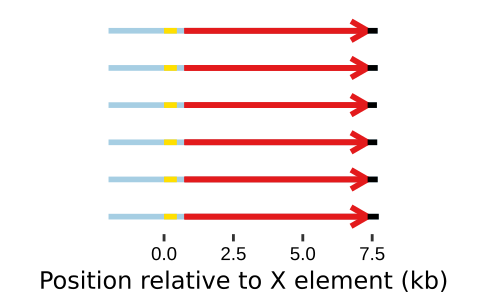

### Chr_V.right.png

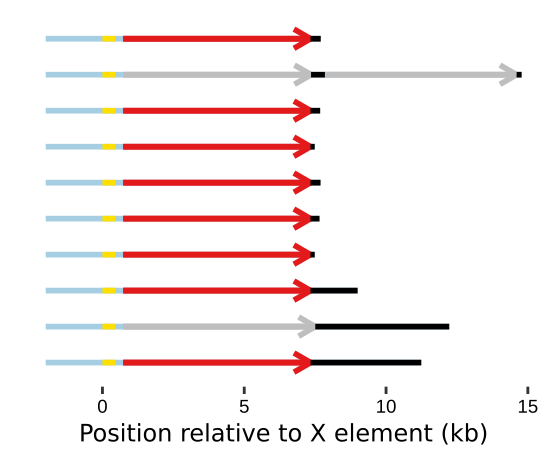

### Chr_VI.left.png

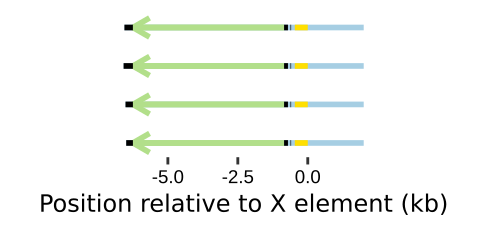

### Chr_VI.left.png

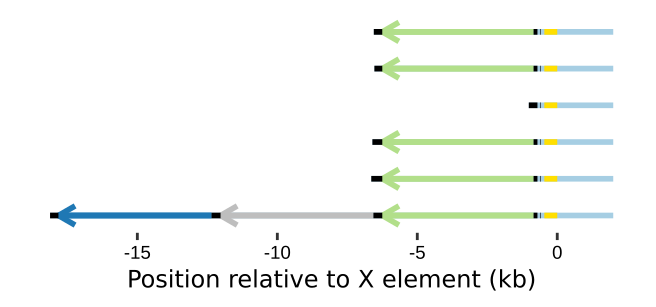

### Chr_VI.left.png

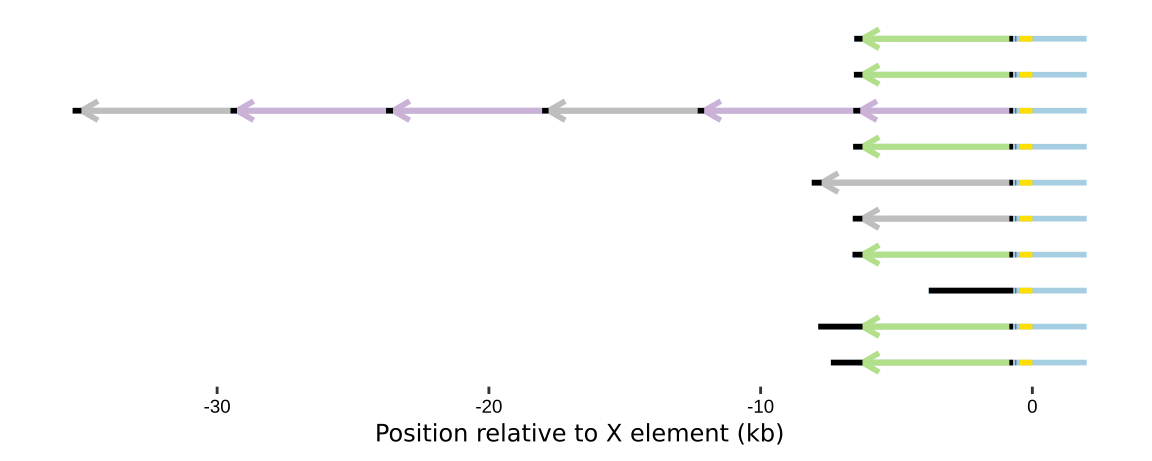

### Chr_VI.right.png

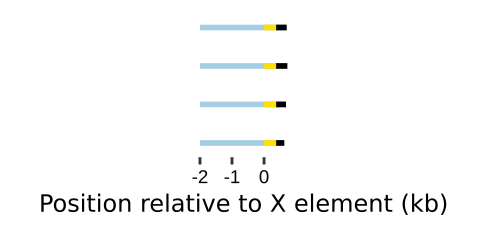

### Chr_VI.right.png

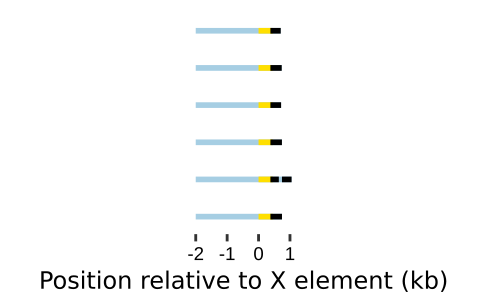

### Chr_VI.right.png

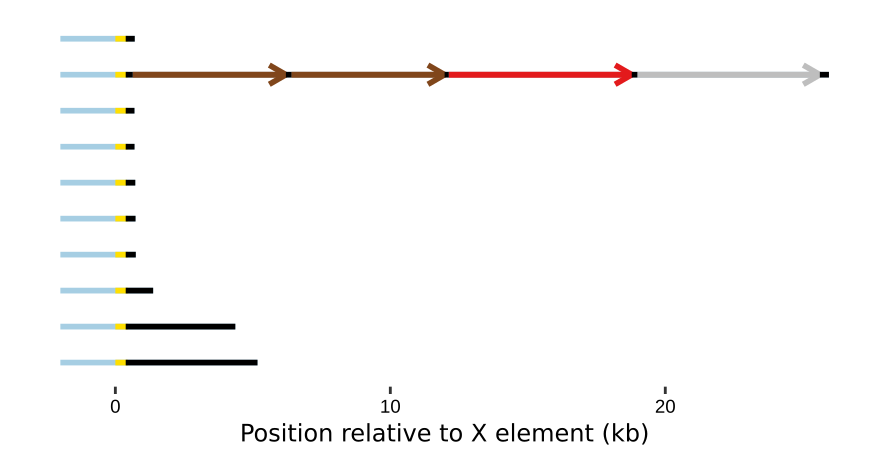

### Chr_VII.left.png

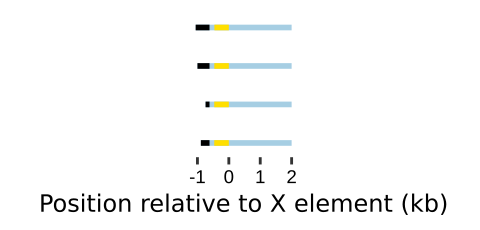

### Chr_VII.left.png

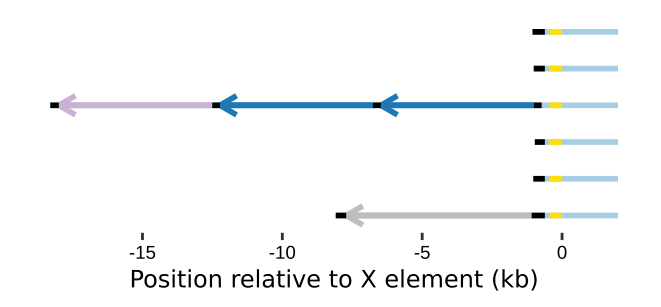

### Chr_VII.left.png

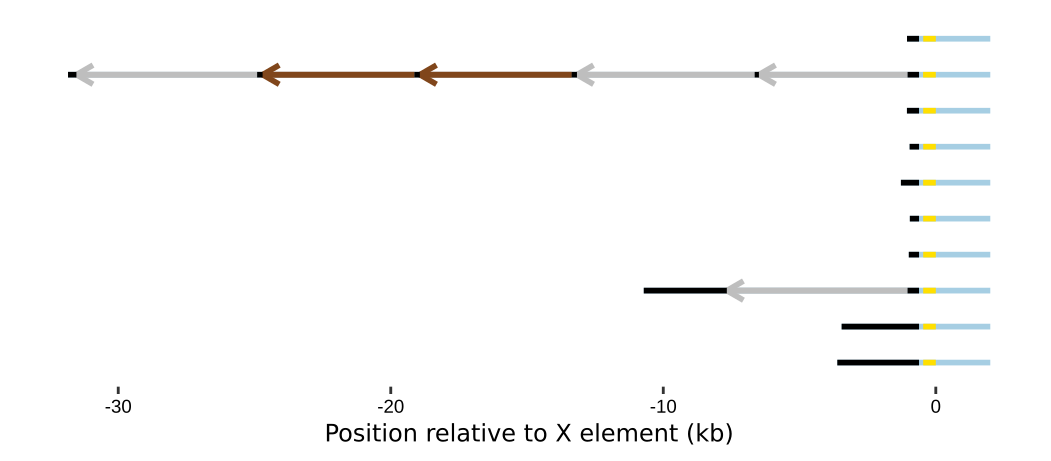

### Chr_VII.right.png

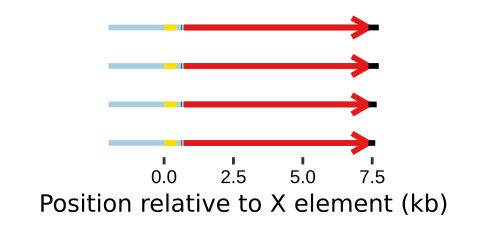

### Chr_VII.right.png

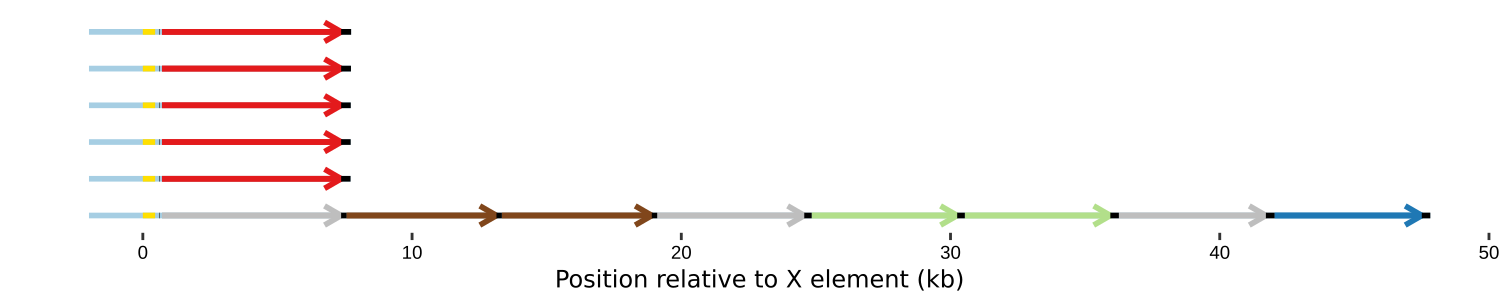

### Chr_VII.right.png

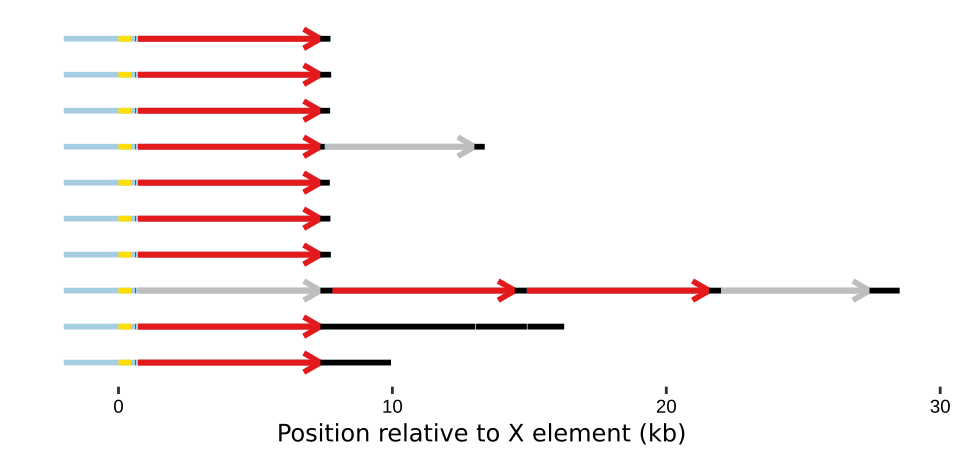

### Chr_VIII.left.png

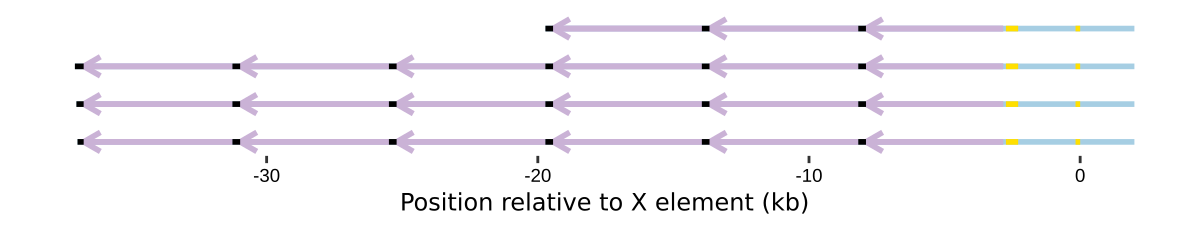

### Chr_VIII.left.png

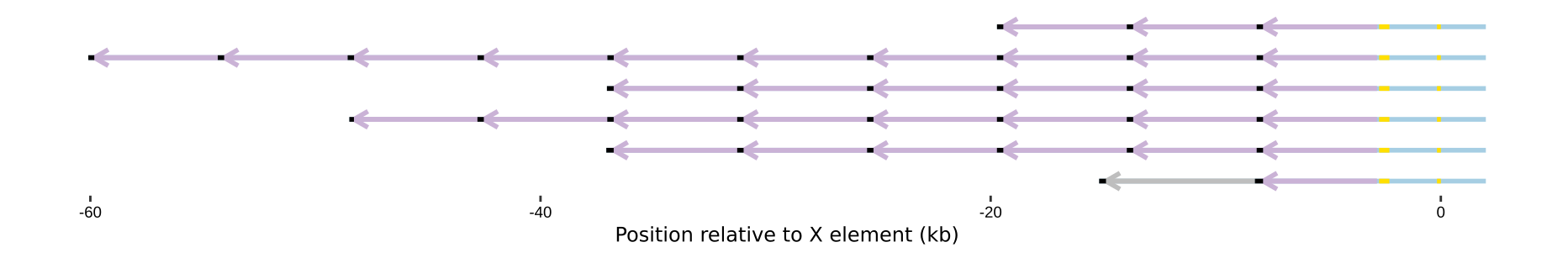

### Chr_VIII.left.png

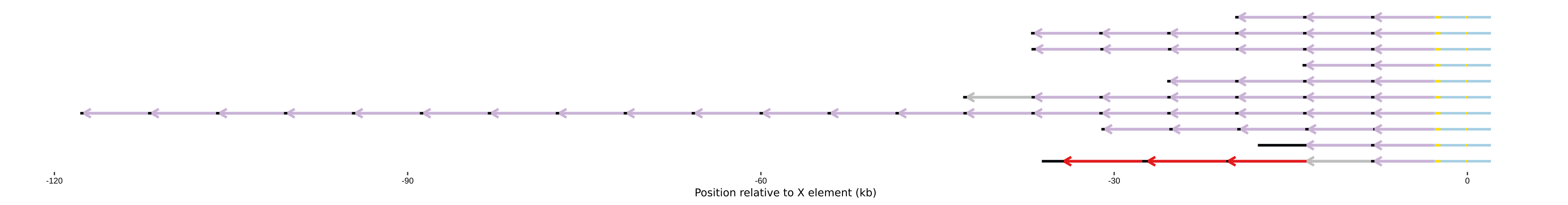

### Chr_VIII.right.png

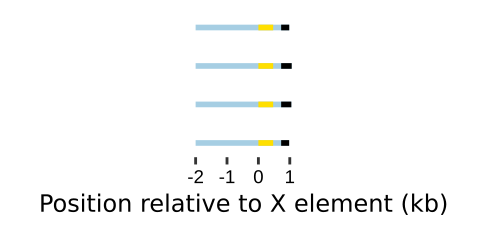

### Chr_VIII.right.png

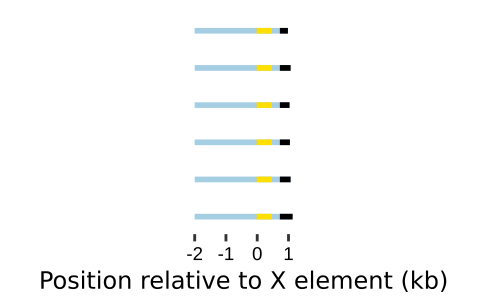

### Chr_VIII.right.png

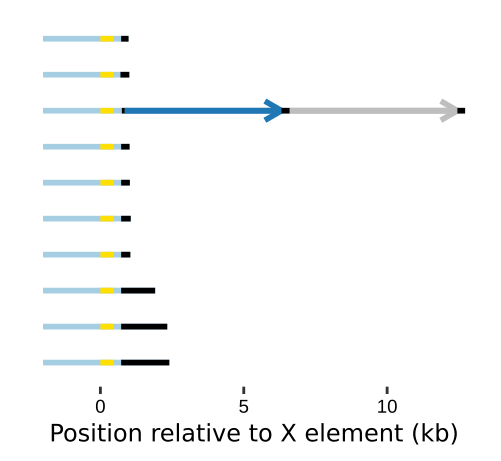

### Chr_X.left.png

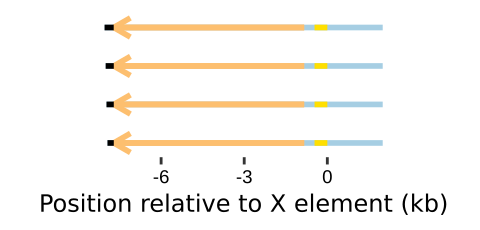

### Chr_X.left.png

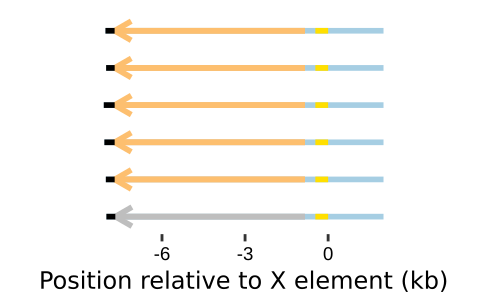

### Chr_X.left.png

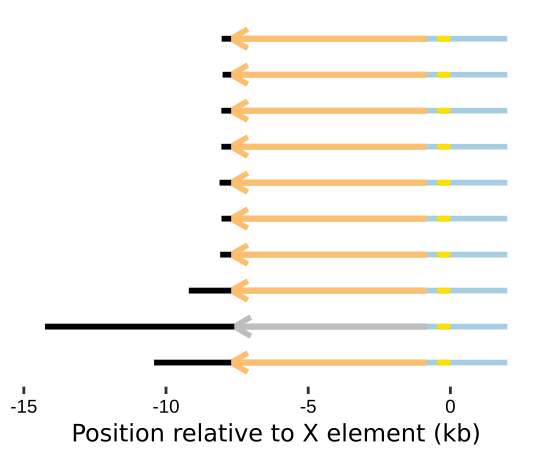

### Chr_X.right.png

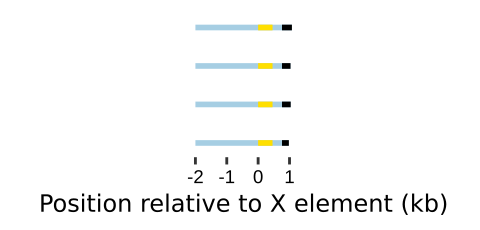

### Chr_X.right.png

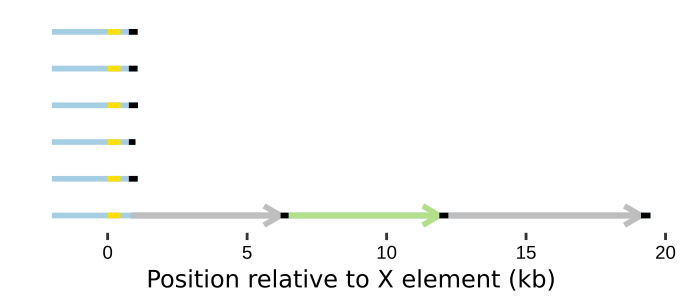

### Chr_X.right.png

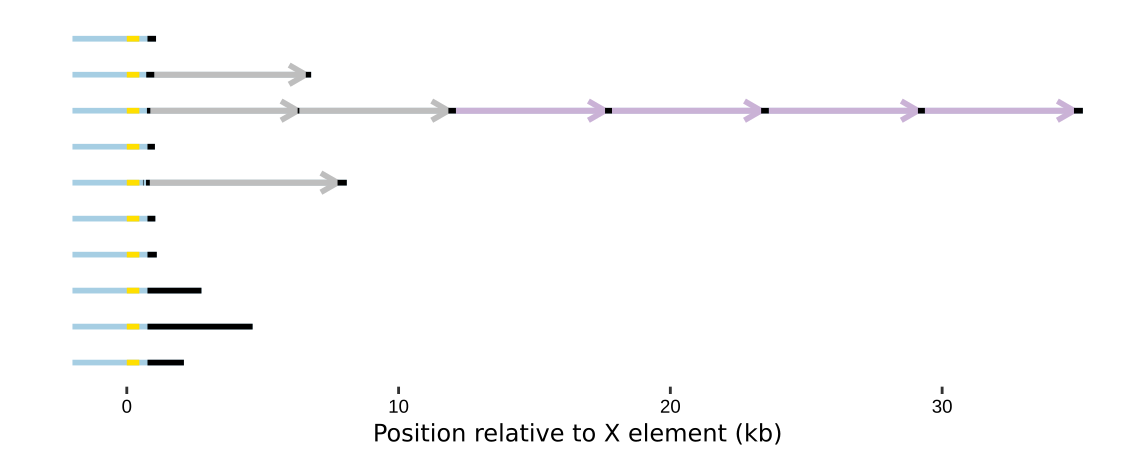

### Chr_XI.left.png

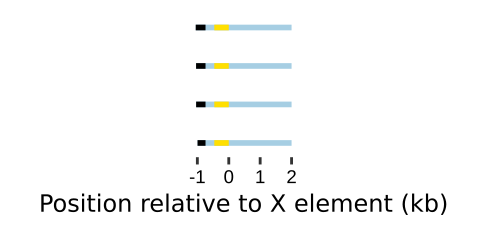

### Chr_XI.left.png

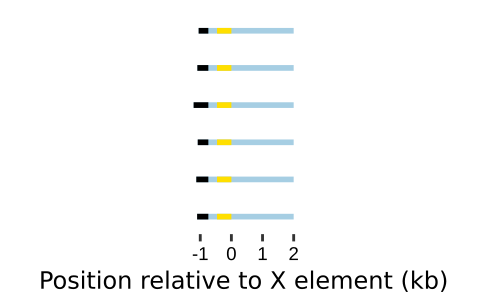

### Chr_XI.left.png

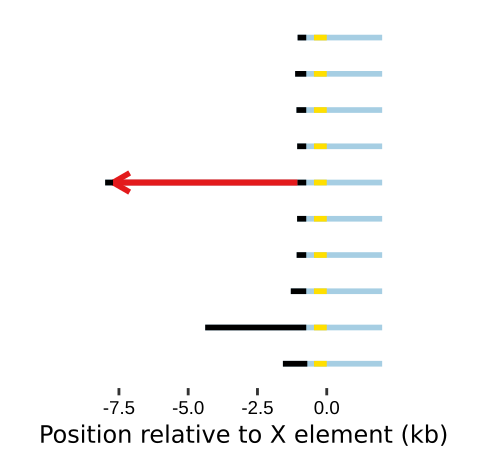

### Chr_XI.right.png

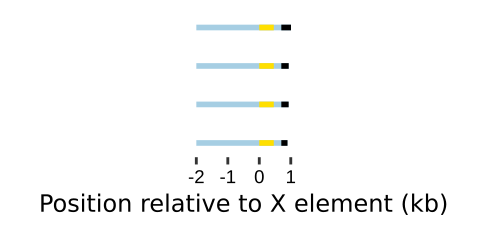

### Chr_XI.right.png

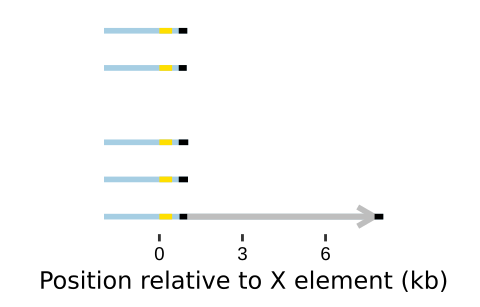

### Chr_XI.right.png

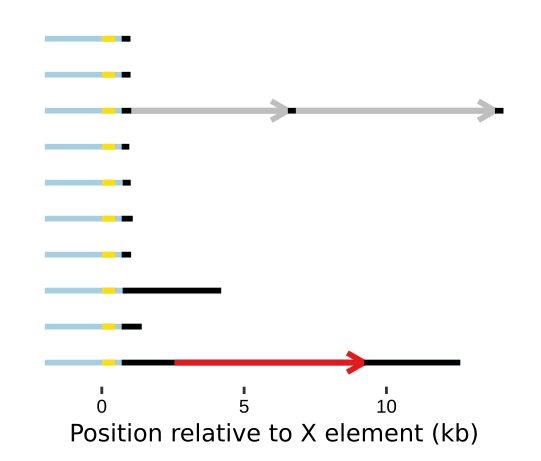

### Chr_XII.left.png

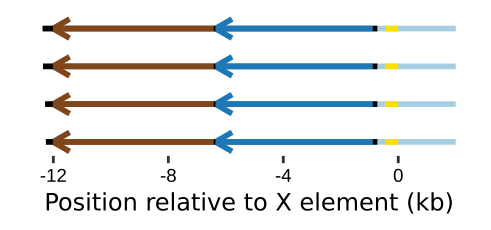

### Chr_XII.left.png

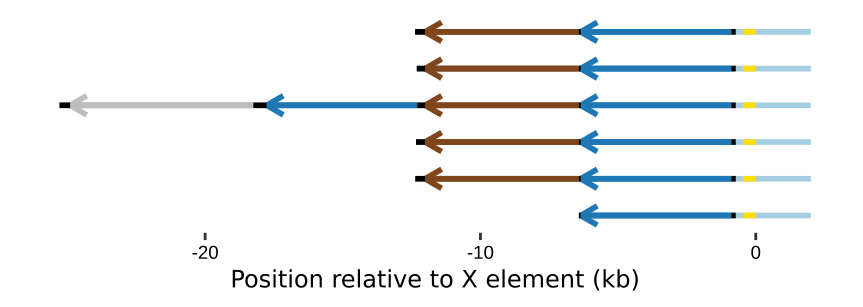

### Chr_XII.left.png

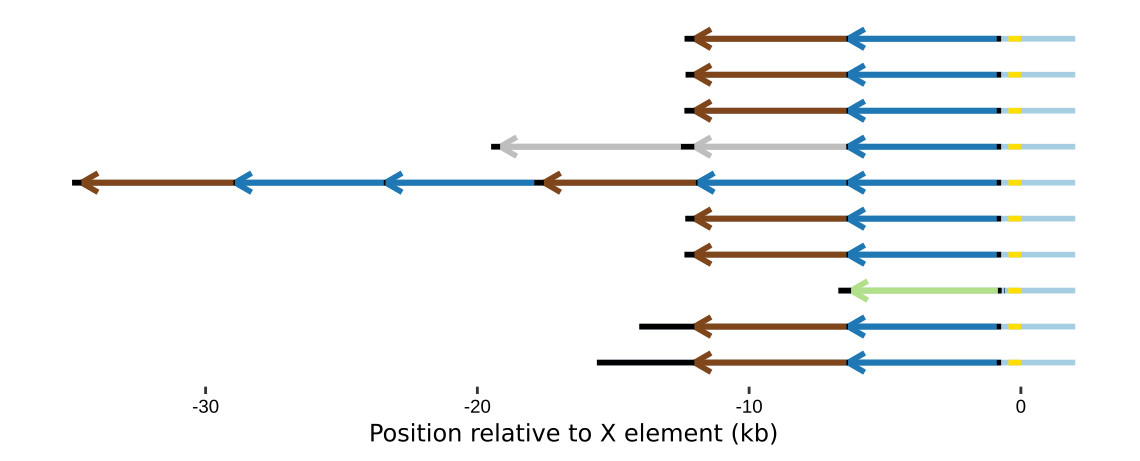

### Chr_XII.right.png

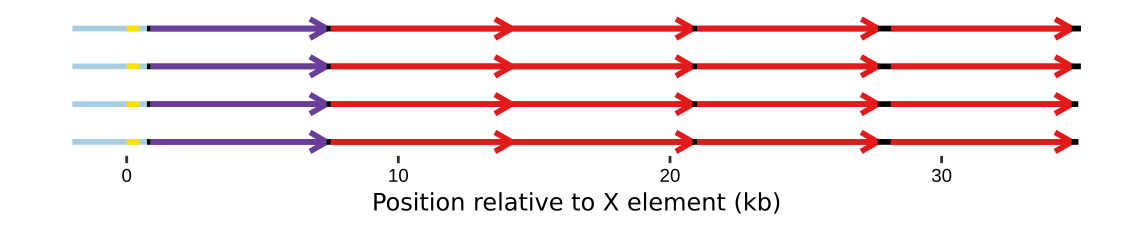

### Chr_XII.right.png

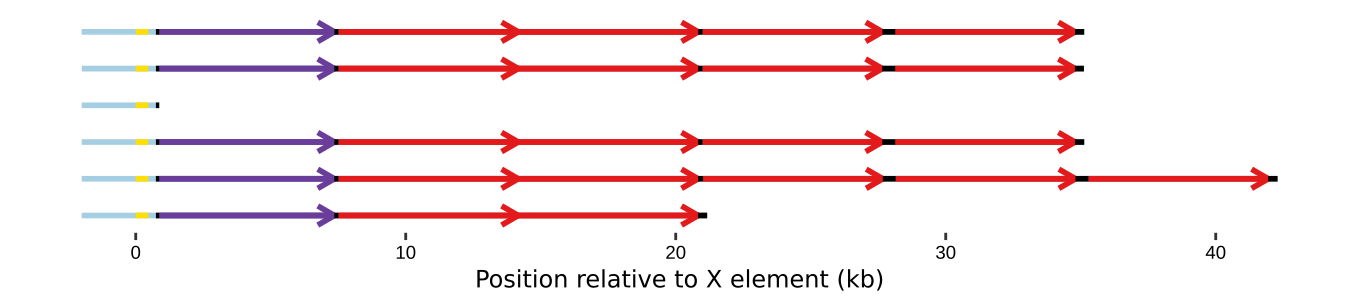

### Chr_XII.right.png

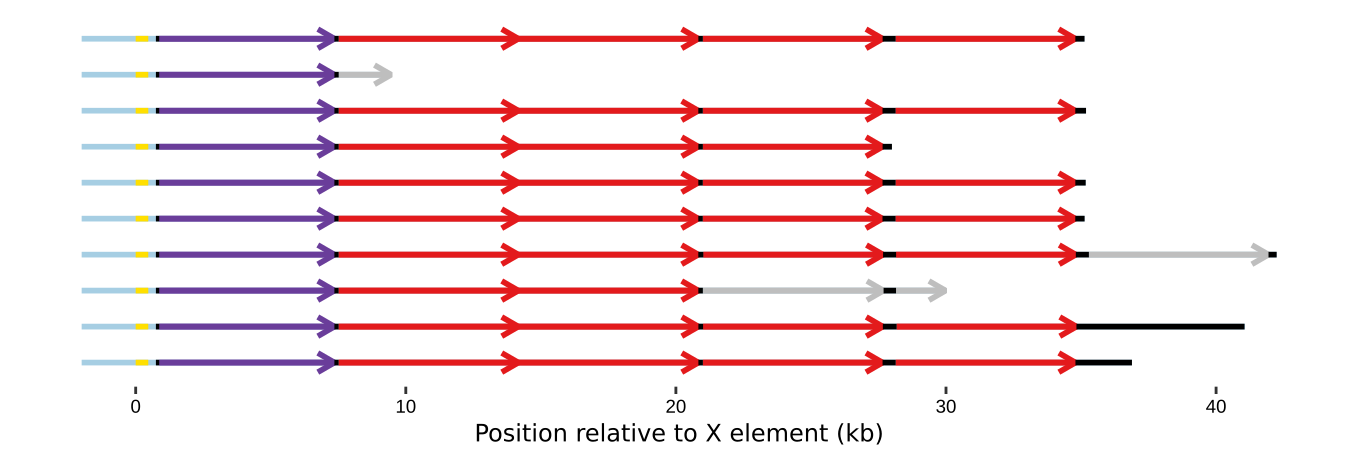

### Chr_XIII.left.png

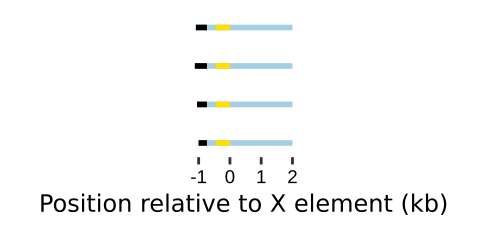

### Chr_XIII.left.png

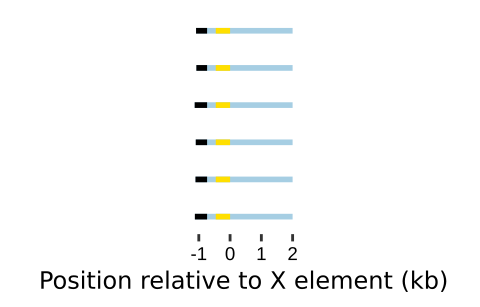

### Chr_XIII.left.png

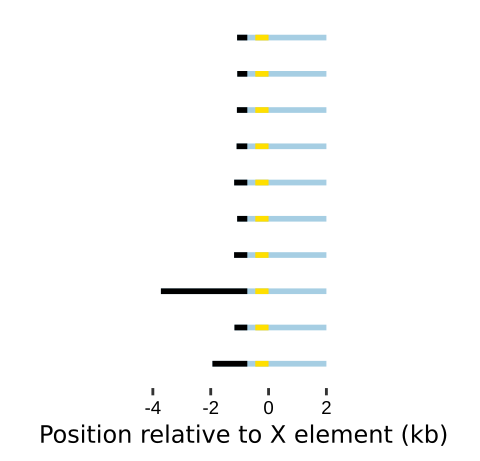

### Chr_XIII.right.png

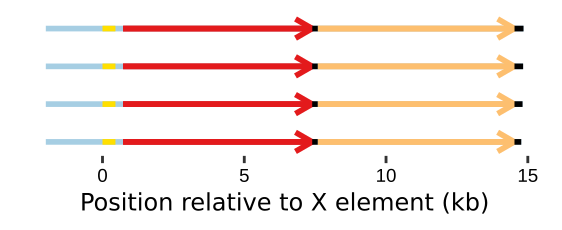

### Chr_XIII.right.png

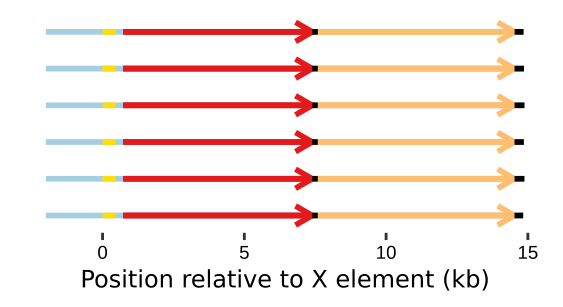

### Chr_XIII.right.png

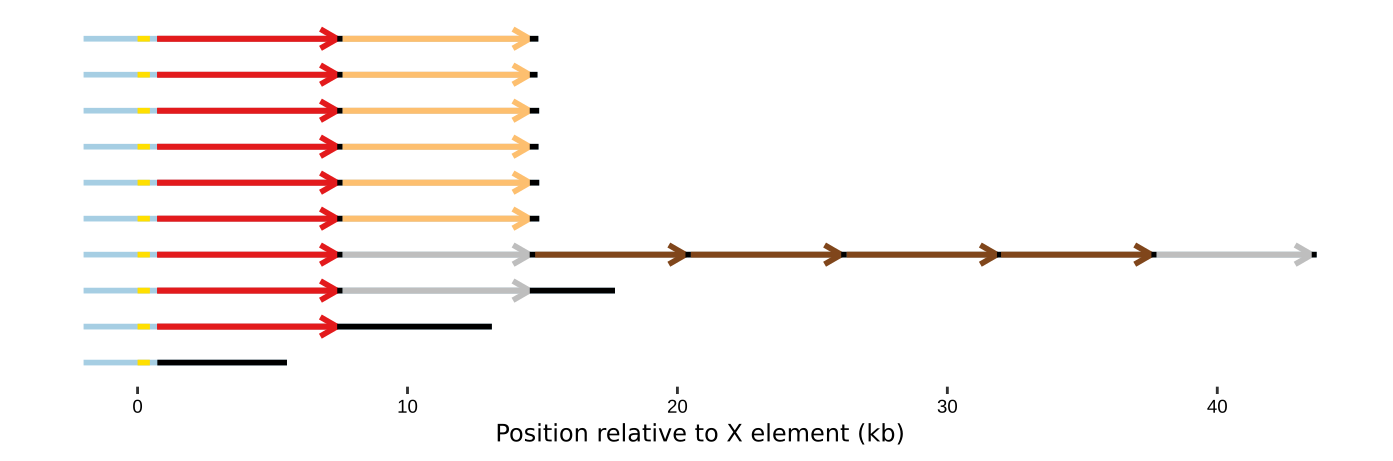

### Chr_XIV.left.png

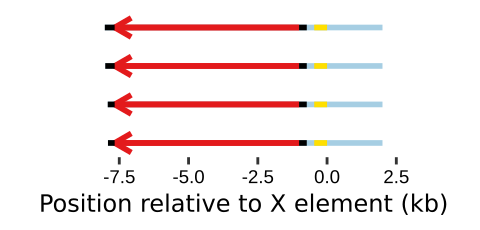

### Chr_XIV.left.png

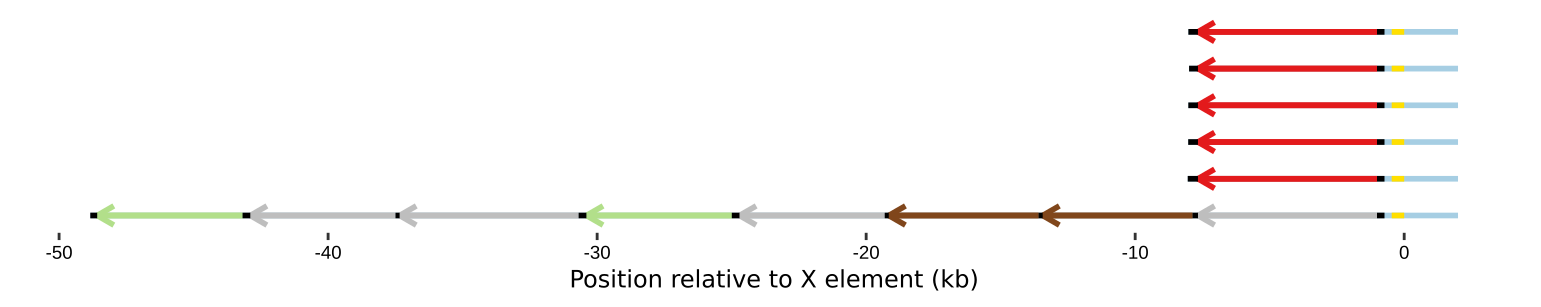

### Chr_XIV.left.png

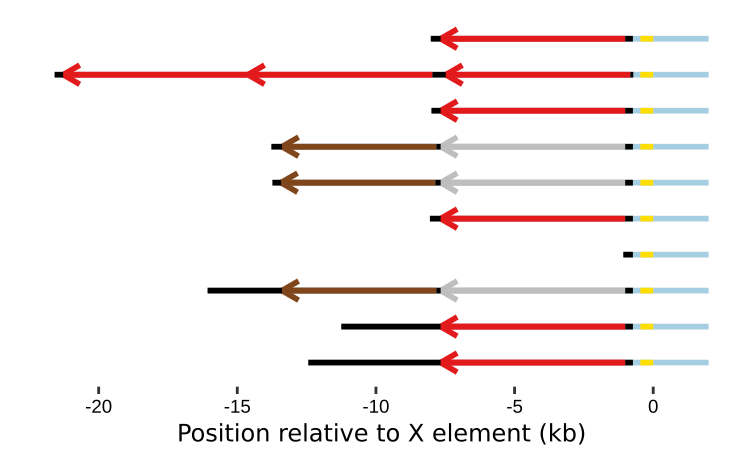

### Chr_XIV.right.png

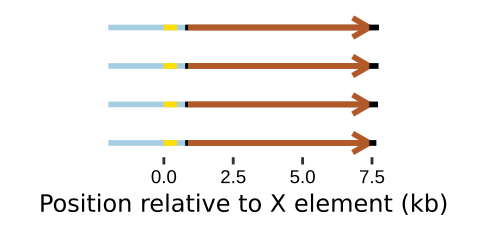

### Chr_XIV.right.png

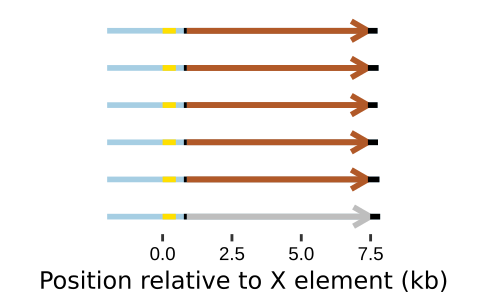

### Chr_XIV.right.png

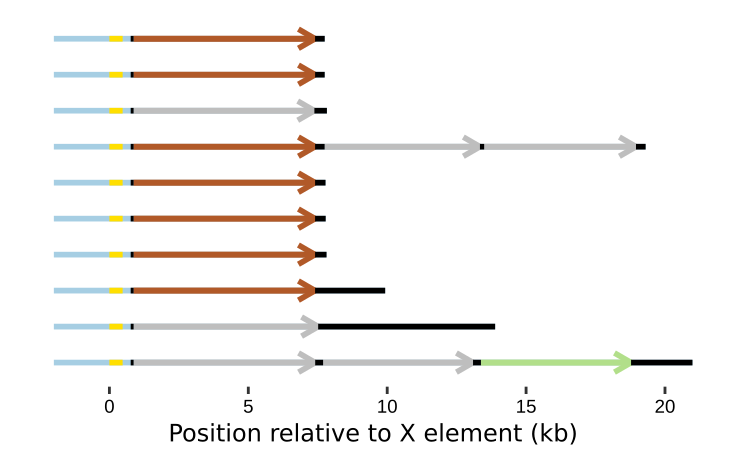

### Chr_XV.left.png

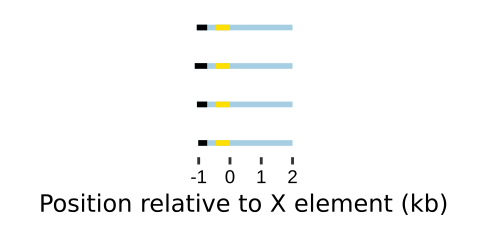

### Chr_XV.left.png

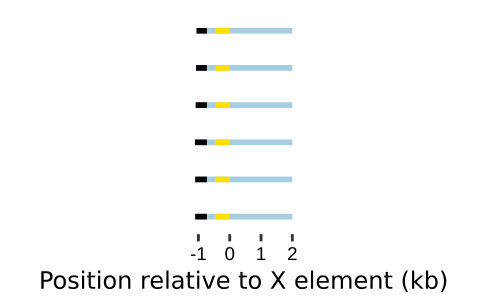

### Chr_XV.left.png

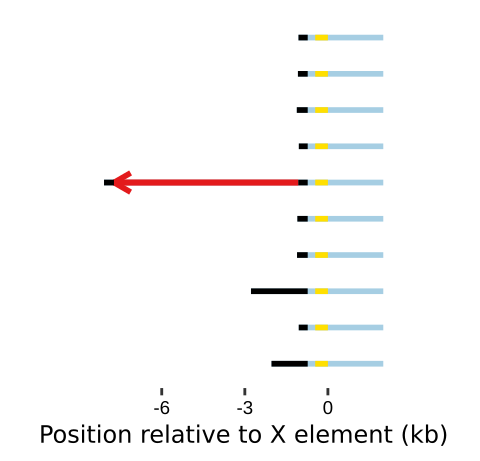

### Chr_XV.right.png

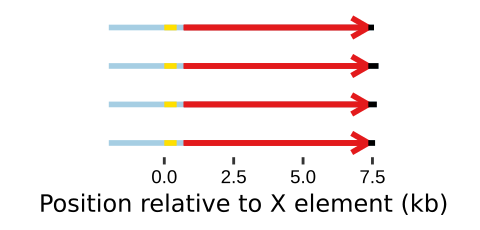

### Chr_XV.right.png

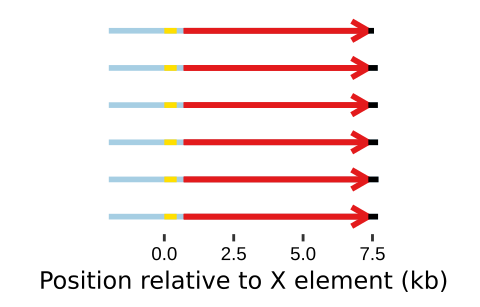

### Chr_XV.right.png

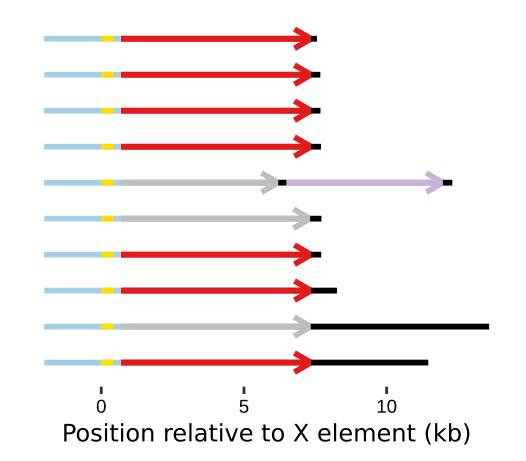

### Chr_XVI.left.png

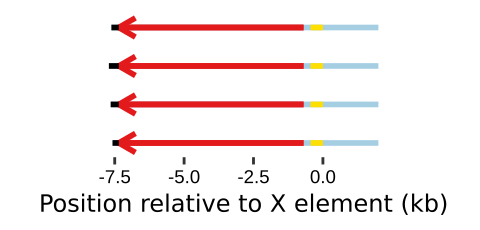

### Chr_XVI.left.png

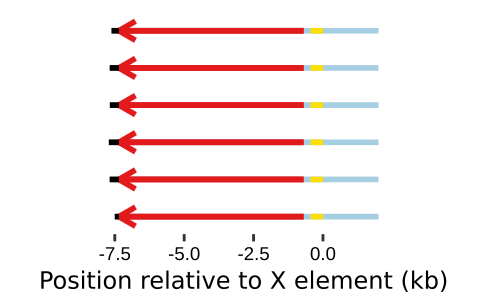

### Chr_XVI.left.png

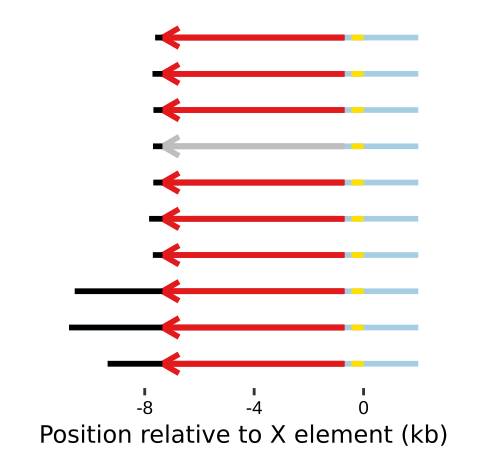

### Chr_XVI.right.png

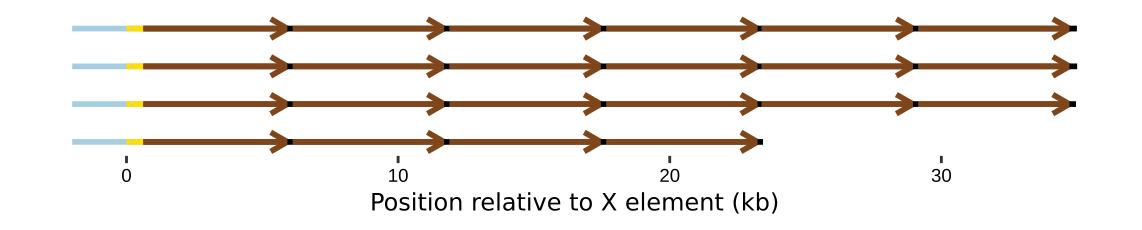

### Chr_XVI.right.png

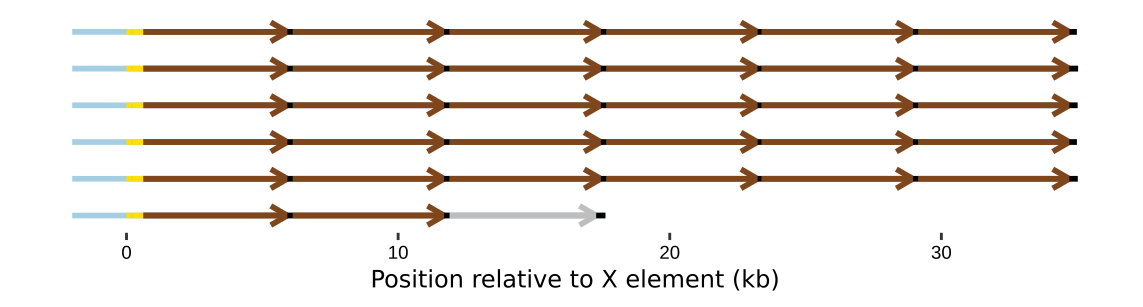

### Chr_XVI.right.png

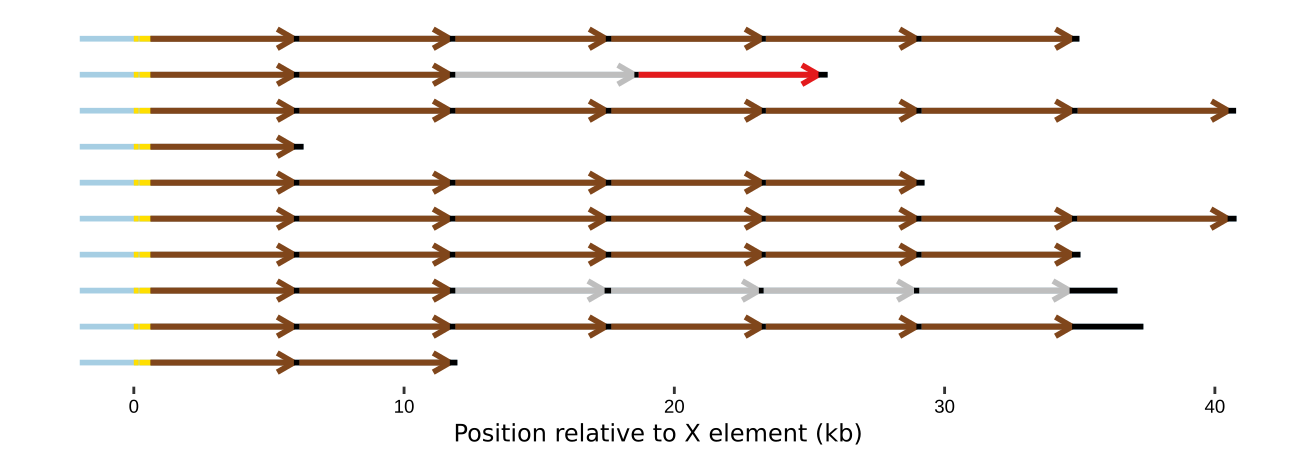
